# Subcellular pharmacology of a novel cell-permeable fluorescent phosphodiesterase inhibitor using advanced fluorescence microscopy

**DOI:** 10.64898/2026.08.22.746441

**Authors:** Soumyabrata Banik, Michele Anselmi, Jagannath Sathpathy, Pier Giorgio Cozzi, Hannes Schihada, Benjamin T. Goult, Andrea Gualandi, Paolo Annibale

## Abstract

Direct visualisation of drug–target engagement within living cells remains a major challenge. Here we develop a cell-permeable fluorogenic analogue of the phosphodiesterase 4 (PDE4) inhibitor rolipram (MAN193), generated by conjugation to fluorescein diacetate. Following intracellular activation, the probe functions as a near-neutral PDE4 antagonist and enables direct visualisation of endogenous PDE4 populations. Using advanced fluorescence spectroscopy imaging approaches, including molecular brightness analysis and fluorescence anisotropy imaging, we demonstrate rolipram-displaceable binding of the analogue to cytosolic PDE4 and resolve thesubcellular distribution of binding sites. We further show that the probe enables visualisation of untagged PDE4 at focal adhesion complexes in cardiomyocyte-like cells. Together these findings establish fluorogenic drug conjugation combined with quantitative spectroscopy imaging as a generalisable strategy to map intracellular drug–target engagement with subcellular resolution, providing an effective framework for interrogating the spatial pharmacology of small molecules in living systems.

## Introduction

Within an organism, the way pharmaceutical compounds (drugs) exert their action is typically associated to the general concepts of potency and efficacy. How well the drug elicits an effect defines the efficacy, how much drug is necessary to achieve the effect is associated to potency (Weatherall 2016). These concepts ultimately depend on a drug reaching a specific cellular target (proteins, nucleic acids), binding to it and remaining bound for a sufficient time in order to elicit an effect.

How much drug will be bound at any given time to a target will depend on the microscopic kinetic rates of the pharmacophore, that in turn crucially depend on its local concentration. If the drug is, for instance, sequestered in the ‘wrong’ cellular compartment, its concentration at the target will drop, reducing binding. This will in turn require a higher dose to be administered, increasing the likelihood of side effects: for instance, the lysosomal sequestration of certain chemotherapeutics is a well-known phenomenon (Hraběta et al. 2020). Conversely, the subcellular targeting of a compound could be the key to its efficacy. A striking example of such phenomenon is provided by the antituberculotic antibiotic bedaquiline, where its main pathway to target m*ycobacterium tuberculosis* is due to its highly lipophilic character which leads its accumulation in intracellular lipid droplets that the bacteria uses as sources of lipids for their metabolism within the host (Greenwood et al. 2019).

It would be thus desirable to visualise individual drug molecules’ interactions within a cell, and indeed reports of the visualisation of endogenous and exogenous pharmacophores using fluorescence date back to the 1960s. Tissue localisation of monoamines, such as dopamine and adrenaline through fluorogenic reactions was reported in 1964 (Carlsson et al. 1964). Taylor and Wang in 1980 reviewed current progress in cellular delivery of fluorescently labelled molecules, highlighting several examples of fluorescent drug conjugates, prominently drugs affecting microtubules (colchicine) and cytoskeletal actin (phalloidin); the latter is nowadays used routinely as an actin stain. Concanavalin A fluorescent conjugates were used to conduct early-stage investigation of ligand-receptor interactions, followed by several applications to other membrane receptors ligands (Taylor and Wang 1980). While the generation of fluorescent analogues of small molecules and peptide receptor ligands has developed significantly in the last decades (Daly and McGrath 2003), the development of fluorescent drug analogues of pharmacophores directed at intracellular targets has proven more challenging, given the complex physicochemical environment the drugs are exposed to after crossing the cell plasma membrane. Examples include fluorescent conjugates of the Src kinase inhibitor dasatinib (Vetter et al. 2014), and PARP inhibitors (Vinegoni et al. 2017), which have been used to monitor intracellular drug uptake and distribution.

However, fluorescence intensity readouts from analogues aimed at intracellular targets suffers from poor signal to background fluorescence, since the unbound ligand cannot be washed off as easily as an extracellular ligand, necessitating alternative approaches, such as the development of probes that become fluorescent upon interaction with target enzymes. Most of these developments are in rather preliminary stages with respect to translation to fluorescent drug analogues of actual pharmacophores (Wysocki and Lavis 2011).

Within the suite of fluorescence spectroscopy approaches available with the pool of already well-established fluorophores, there are tools that can be brought to bear towards the goal of visualising the subcellular dynamics of fluorescently-tagged pharmaceutical drugs. Amongst these are (i) image fluctuation spectroscopy approaches, that can monitor changes in oligomerisation, a fingerprint of a fluorescent drug analogue going from a monodisperse, cytosolic free state, to a state bound to its target, that are unlikely to be diffuse and monodisperse (Işbilir et al. 2021); (ii) fluorescence anisotropy approaches, that have been used to conduct binding assays ‘in-cuvette’ for decades (Jameson and Ross 2010), and have more recently being applied again to the problem of generating images of intracellular fluorescent drug analogues (Dubach et al. 2014).

Here we develop a strategy that combines fluorogenic drug conjugation with fluorescence spectroscopy imaging to visualise drug–target engagement with subcellular resolution in living cells. Our approach exploits fluorescein diacetate (FDA), a membrane-permeable non-fluorescent precursor that is converted intracellularly into emissive fluorescein by endogenous esterases (Rotman and Papermaster 1966). By coupling FDA to pharmacophores, we generate probes that become fluorescent selectively within the intracellular environment while remaining compatible with a broad range of spectroscopic imaging approaches. As a proof of principle, we developed a fluorescent drug analogue of rolipram, a Phosphodiesterase 4 (PDE4) inhibitor (Paes et al. 2021). PDE4 plays a central role in regulating intracellular cAMP levels required for the specificity of selective G protein-coupled receptors (GPCR)-mediated extracellular stimuli thanks to its ability to establish intracellular cAMP gradients (Bock et al. 2020). We show that the rolipram-FDA is efficiently taken up by cells, converted into intracellular rolipram-fluorescein and binds specifically to cytosolic PDE4s. Using advanced fluorescence spectroscopy imaging, we further demonstrate visualisation of endogenous, untagged PDE4 and reveals its localisation at talin-1-containing integrin adhesion complexes in cardiomyocyte-like cells.

## Results

We set out to generate a fluorescent analogue of the well-established PDE4 inhibitor, rolipram (**Figure 1**). To achieve this, we adapted a strategy previously used to fluorescently label cAMP (Bock et al. 2020), employing fluorescein diacetate (FDA) as a cell-permeable, fluorogenic moiety.

**Figure 1.**
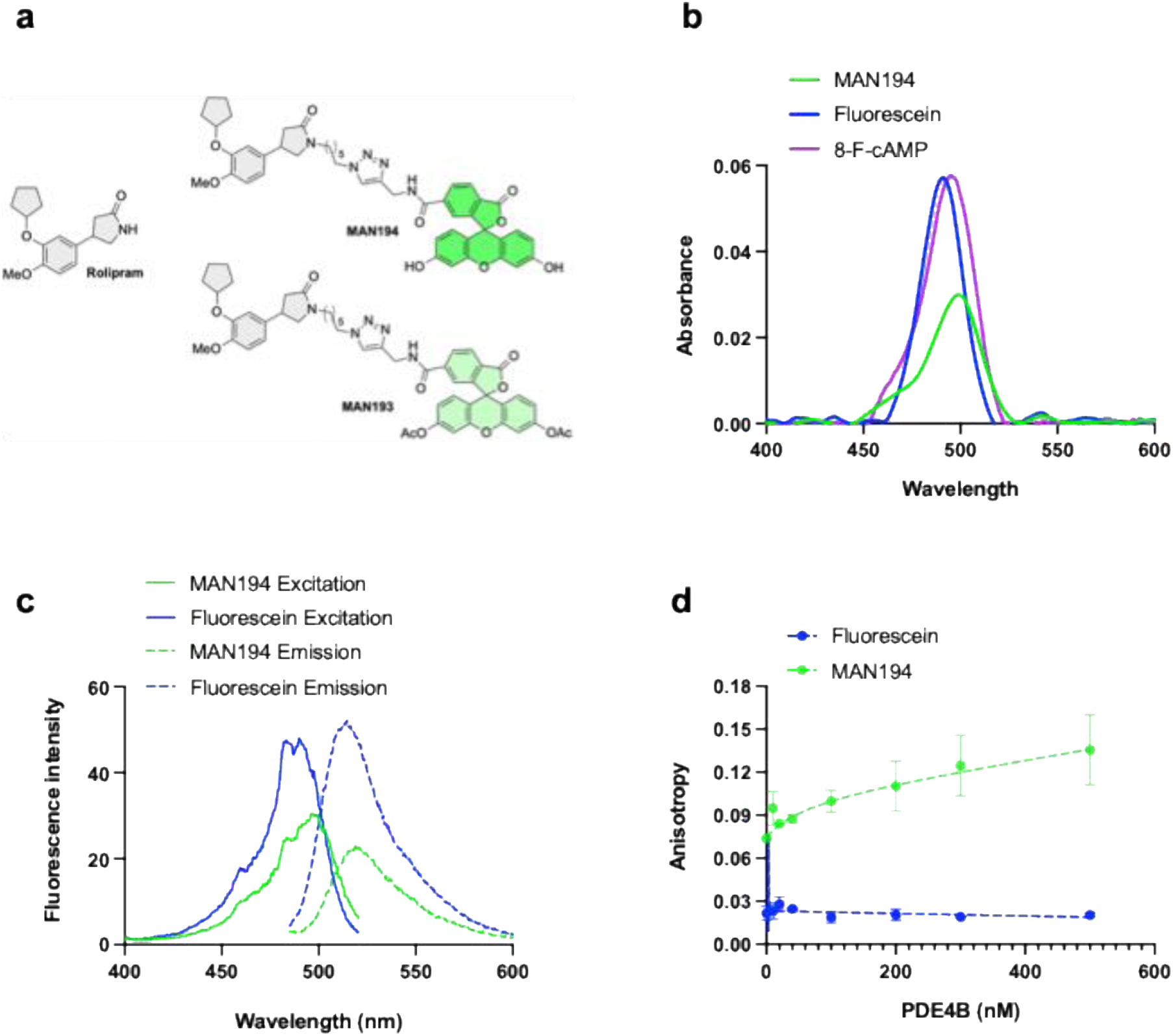
Synthesis and characterisation of MAN193 and MAN194. a) Chemical structures of rolipram, the fluorogenic precursor MAN193 generated by conjugation to fluorescein diacetate (FDA), and the intracellularly de-esterified fluorescent form MAN194. b) Absorption spectra of MAN194, fluorescein and 8-F-cAMP. c) Excitation (solid lines) and emission (dashed lines) spectra of MAN194 and fluorescein. Emission maxima were observed at 520 nm for MAN194 and 514 nm for fluorescein. d) Binding isotherm of MAN194 to purified PDE4B catalytic domain. Drug-protein binding assay based on fluorescence anisotropy recorded using a Cary Eclipse fluorescence spectrophotometer (Varian). With 460 nm as excitation wavelength, and emission recorded at 520 nm, fluorescence anisotropy was recorded by titrating 100 nM MAN193 against increasing concentration of PDE4B. Binding curves were fitted using GraphPad prism. Data represent mean ± s.e.m. from n=2 independent experiments.

Guided by molecular docking studies (Dym et al. 2002) and reported structures of PDE4 bound to rolipram (Cedervall et al. 2015), FDA was conjugated to the rolipram pharmacophore such that the fluorophore was positioned distal to the PDE4-binding warhead.Synthesis was accomplished through a copper-catalysed azide–alkyne cycloaddition (CuAAC, “click” chemistry) between an azido-functionalised rolipram derivative bearing a C6 linker at the amide nitrogen and an alkyne-functionalised FDA derivative, prepared as the corresponding propargyl amide. The synthetic procedures are described in **Appendix 1**. The resulting compound, designated MAN193 prior to deacetylation, and its deacetylated fluorescein form MAN194 are shown in **Figure 1a**. MAN194 displayed absorbance spectra broadly consistent with that of fluorescein, albeit with a modest red-shift and reduced extinction coefficient (**Figure 1b**). A similar redshift is observed in the excitation and emission spectra (**Figure 1c**).

To determine whether MAN194 bretained PDE4 binding, we used an *in vitro* anisotropy-based binding assay with purified PDE4B catalytic domain. As MAN194 binds to PDE4B, its rotational diffusion reduces, and therefore the measured steady state fluorescence anisotropy (*r*) increases. The steady state anisotropy for free MAN194 in solution was r_0_ = 0.075 ± 0.003, compared to fluorescein 0.033 ± 0.008**. Supplementary Figure 1a** further illustrates that MAN194 displays a different behavior when compared to fluorescein in a Perrin plot. We measured the fluorescence lifetime of MAN194 and fluorescein, observing a lifetime *τ* = 4.01 ns for fluorescein -in line with reported values (ISS) - and a double-exponential lifetime for MAN194 with values of *τ*_1_ = 3.41 ns and *τ*_2_ = 1.51 ns (**Supplementary Figure 1c-g**). We include here also values from the previously reported 8-F-cAMP as an example of another analogue originating from FDA fusion (Bock et al. 2020). Together, these data indicate that both increased rotational diffusion time (due to the larger molecular weight), and reduced fluorescence lifetime contribute to the altered slope of MAN194 in the Perrin plot.

**Supplementary Figure 1.**
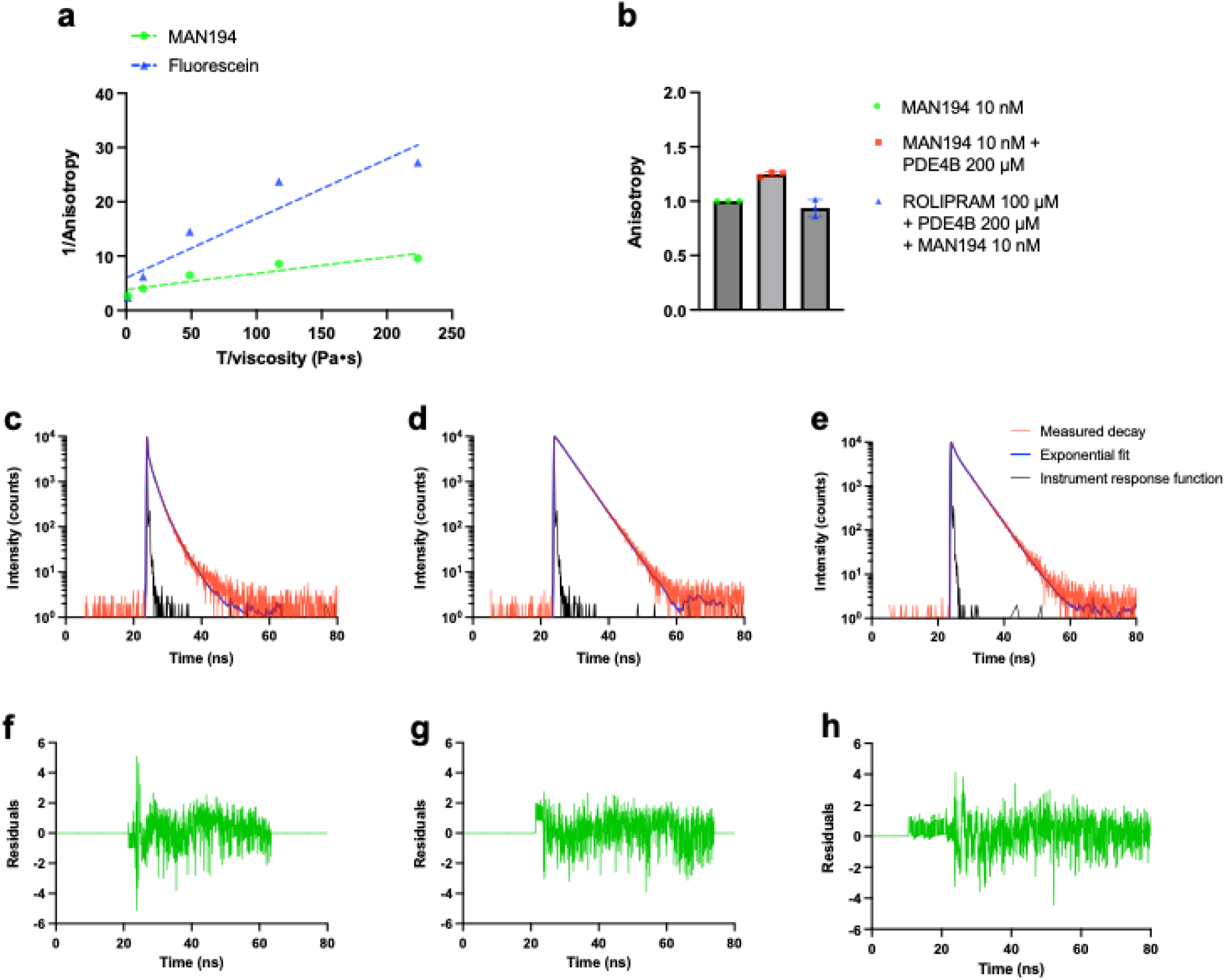
Fluorescence anisotropy and lifetime characterisation of MAN194. Biophysical and spectral validation of the probe. a) Instrument validation to differentiate anisotropy measurements based on the rotational diffusion of molecules. Anisotropy values of 100 nM MAN194 and Fluorescein dissolved in 10-90% (w/w) water-glycerol solutions are plotted as 1/anisotropy against T/viscosity. The data was fitted using simple linear regression using GraphPad Prism. b) specific binding of fluorescently labelled drug realised by measuring anisotropy of drug alone, drug with PDE4B catalytic subunit protein and by pre-blocking the protein with 100 µM rolipram before treating with the fluorescent drug (10 nM). Results are plotted as bar chart (mean ± s.e.m.) with each dot representing an individual experiments (n=3 independent experiments). Fluorescence lifetime measured for c) MAN194, d) Fluorescein and e) 8-F-cAMP. The black lines represent the instrument response function (IRF), the red is the observed fluorescent decay curve from the TCSPC experiment, and the blue is for the exponential fit. For the fluorescein and 8-F-cAMP molecule, the data is de-convoluted with the IRF and then fitted with the double exponential decay fit function as shown by the blue line. MAN194 demonstrated two lifetimes of 1.51 ns and 3.41 ns. Similarly, the fluorescence lifetime obtained for fluorescein and 8-F-cAMP are 4.01 ns and 4.08 ns respectively. (f-h) the residuals for the double exponential fitted function (in green) for MAN 194, Fluorescein and 8-F-cAMP respectively.

By titrating PDE4B against a fixed concentration of MAN194, it was possible to build a binding isotherm (**Figure 1d),** showing nanomolar affinity binding of MAN194 to PDE4B (K_d_= 61 nM). This affinity is reduced relative to reported values for rolipram binding, including∼20 nM for [3H]-rolipram binding to rat cerebral cortical membrane preparations (Zhao, Zhang, and O’Donnell 2003), and of ∼1.5 nM for for purified PDE4B (Rocque et al. 1997).

Binding of MAN194 was specific to the PDE4B rolipram-binding site, as preincubation with excess unlabelled rolipram (100 µM) reduced the anisotropy of 10 nM MAN194 to levels comparable to free probe (**Supplementary Figure 1b**).

Together, these data demonstrate that the fluorescent analogue retains the core spectral properties of fluorescein and binds specifically, albeit with a reduced K_d_, to the molecular target of the original pharmacophore.

We next set to monitor the ability of MAN193 to inhibit PDE4 catalytic activity in living cells. We followed a strategy based on bioluminescence sensing of cellular cAMP and conducted concentration-response curves adding our compound to HEK293 cells transiently expressing the GloSensor as well as PDE4B. As cellular cAMP increases, luminescence from the Firefly luciferase increases (Fan et al. 2008; Binkowski et al. 2011). The assay is particularly well-suited o study the addition of fluorescent drug analogues, because it does not require optical excitation, thereby avoiding interference from probe fluorescence. We compared the response of cellular cAMP to addition of increasing concentrations of MAN193 and compared that to the responses to IBMX and rolipram (**Figure 2a,b**). These assays provide us with an IC_50_ value, indicating the ability of the compounds to block 50% of the catalytic activity of cellular PDE4s. We note that conjugation to FDA reduces aqueous solubility relative to rolipram, and therefore the nominal extracellular concentrations reported may underestimate intracellular accumulation of the compound. The data were collected as time series (**Supplementary Figure 2**) and the GloSensor response after about 16 minutes (t = 960 s) was used to generate concentration response curves. IBMX addition, resulted in an inhibition of PDE4 enzymatic activity with an IC50 of 5 ± 2 µM (**Figure 2c**). Rolipram, displayed an IC50 of 0.4 ± 0.4 µM and MAN193 an IC50 of 13 ± 5 µM. The reduced potency of MAN193 is consistent with the lower binding affinity observed in vitro (**Figure 1d**) relative to reported values for rolipram.

**Figure 2.**
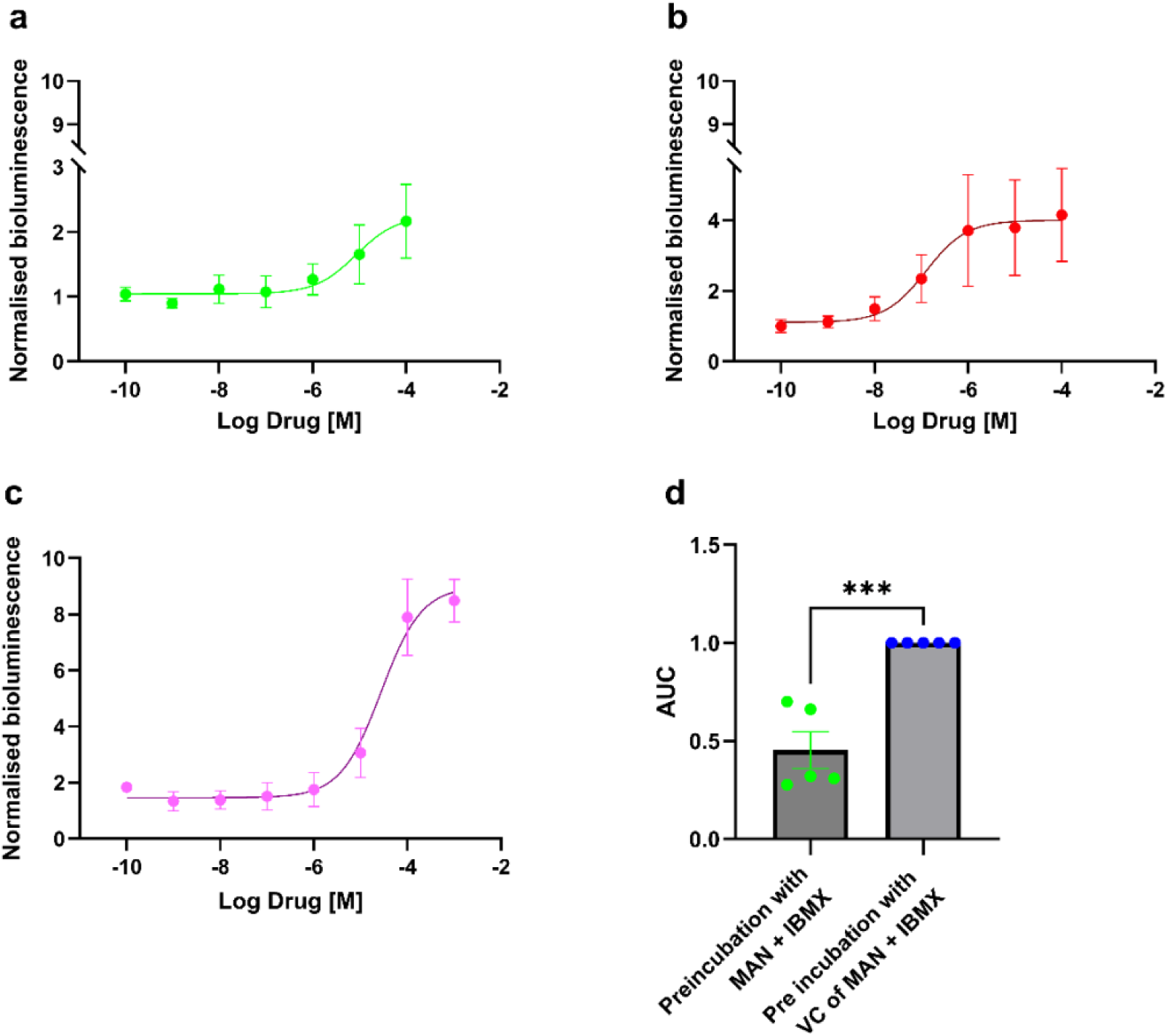
Pharmacological characterisation of MAN193 in PDE4B-overexpressing HEK293 cells. a-c) Concentration-response curves for cAMP levels measured using the GloSensor cAMP assay in HEK293 cells overexpressing PDE4B following treatment with a) MAN193, b) rolipram and c) IBMX. Measurements were recorded as kinetics of 25 cycles with cycle time of 132 seconds. Luminescence signals were normalised to vehicle control. The concentration response curve was generated from the time point at 960 s after ligand addition. IC_50_ values were acquired by fitting the data with Nonlinear fit function in GraphPad Prism using Hill’s equation (Hill slope = 1). The IC_50_ values for MAN193, rolipram and IBMX are 13 ± 5 µM, 0.4 ± 0.4 µM and 5 ± 2 µM respectively whereas E_max_ values are 2.2 ± 0.1, 3.9 ± 0.1 and 9.9 ± 0.4 respectively. Data are mean ± s.e.m. from n=3 independent experiments. d) Blocking experiment performed by pre-treating cells with MAN193 or its vehicle control, followed by measurement of luminescence upon addition of IBMX. Results represented as dot plots with mean ± s.e.m. Unpaired t-test was performed as statistical analysis between the groups. Data is from n=5 independent experiments.

**Supplementary Figure 2.**
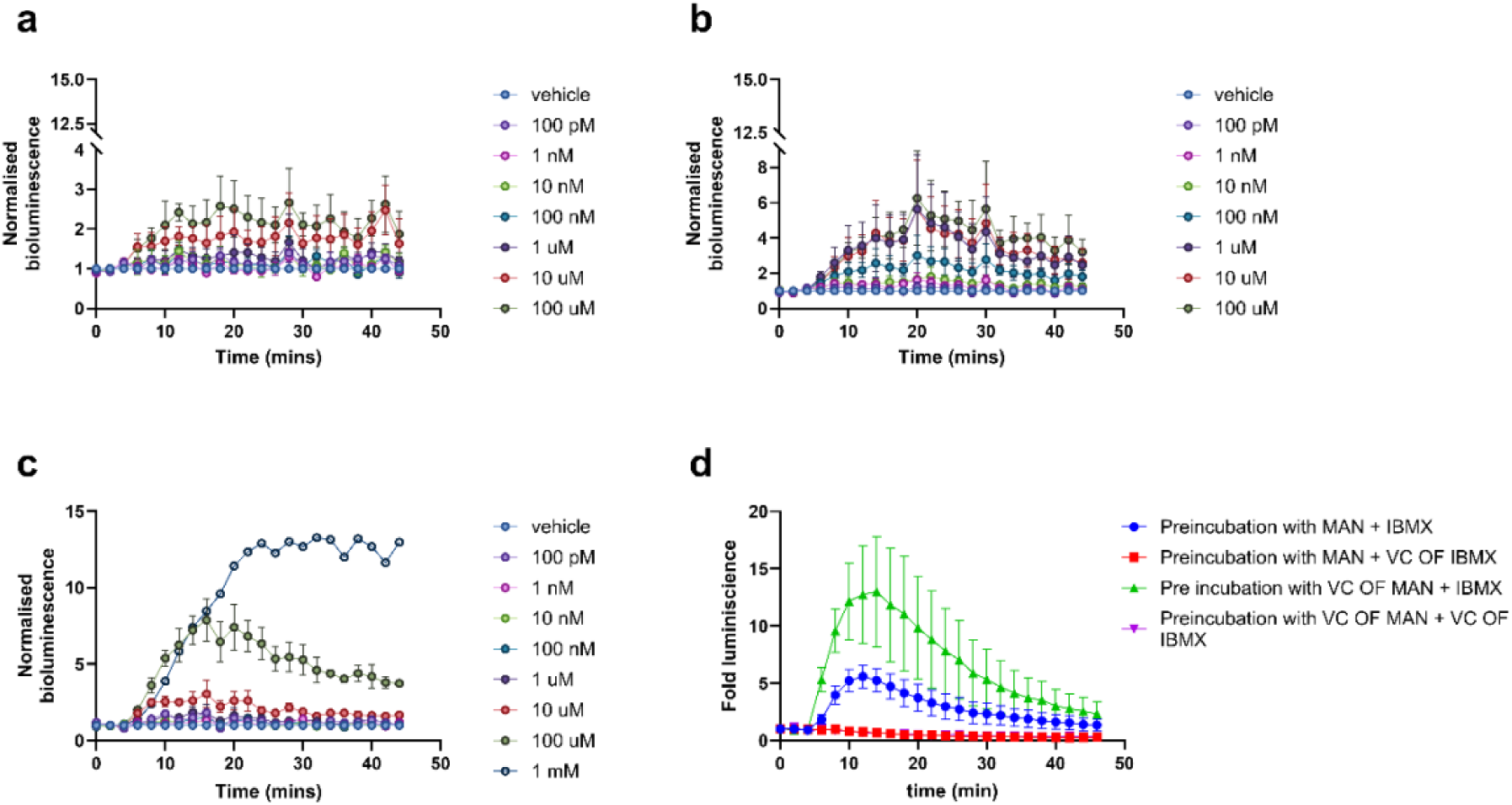
Kinetic response of GloSensor assay. a–c) Time-dependent changes in intracellular cAMP levels following addition of increasing concentrations of a) MAN193, b) rolipram, or c) IBMX, measured using the GloSensor assay in HEK293 cells. Drug concentrations are indicated in the respective panels. d) Time course of cAMP production used to assess the dissociation response.

Our results point also to a reduction in E_max_ of MAN193 relative to rolipram (**Figure 2a, b**), suggesting that this compound functions as a partial PDE4 inhibitor. This is advantageous since for the purpose of non-perturbative visualisation of a target’s localisation, a compound operating as a weak inverse agonist (i.e. a compound reducing basal enzyme activity) or neutral antagonist (i.e. a compound which has no effect on catalytic activity but blocks binding of agonists / stimulators) would be preferred. We confirmed that MAN194 binds to the PDE4 rolipram-binding site in living cells, by preincubating cells with either vehicle or MAN193 prior to adding IBMX. Vehicle-treated cells displayed a significantly greater increase in intracellular cAMP levels compared MAN193-pretreated cells (**Figure 2d**), consistent with competitive occupation of PDE4 binding sites by the fluorescent analogue.

We next addressed the key question of the subcellular distribution of MAN193 in living cells. Although PDE4 inhibition is relevant in a broad range of physiological and pathological conditions, we focus here on its role in compartmentalised signalling. The localisation of PDE4 and subtypes in cells has been the focus of increasing attention, as they play a major role in the compartmentalisation of cAMP, that is in turn required for appropriate cellular response to specific extracellular cues. PDE4 regulates local phosphorylation of signalling proteins, ranging from mechanosensitive proteins to those involved in excitation-contraction coupling in cardiac myocytes. We therefore used both HEK293 cells and the cardiomyocyte-like H9c2 cell line, originating from rat myoblast as model systems to investigate intracellular localisation of the fluorescent analogue.

Confocal time-lapse imaging of H9c2 cells following addition of 10 µM MAN193 revealed progressive intracellular accumulation of fluorescence over time (**Figure 3a**). Because MAN193 becomes fluorescent only following intracellular esterase-mediated de-esterification, the observed signal reflects cellular uptake and intracellular conversion of the probe. Fluorescence intensity increased throughout the cell and reached an apparent steady state within approximately 30 min, indicating efficient intracellular delivery and retention of the fluorescent analogue.

**Figure 3.**
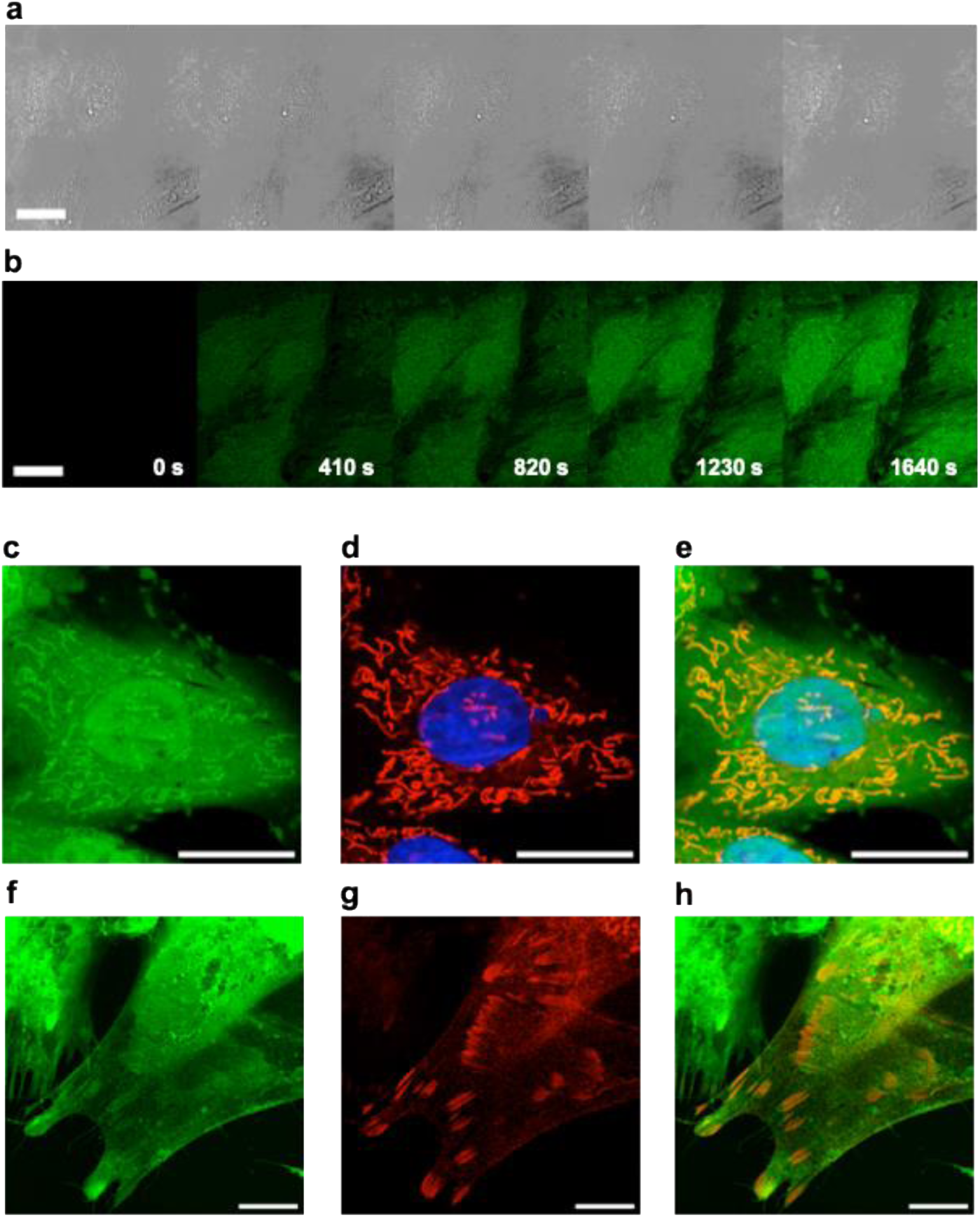
Live-cell uptake of MAN193 and localisation of MAN194 in H9c2 cells. a-b) Time course upon addition of 10 µM MAN193 and intracellular conversion into MAN194 within H9c2 cells, monitored using confocal microscopy (100 Hz linescan rate, 100 nm pixel size, 488 nm excitation, 500-600 nm detection window). Frames shown in the mosaic are acquired every 410 s. c) Cells imaged after 30 min incubation with 10 µM MAN193, further incubated with d) MitoTracker Deep Red (633 nm excitation, 640-700 nm detection window) and Hoechst (405 nm excitation, 430-470 nm detection window). e) Merge of all colour channels. f) Cells imaged following 30 min incubation with 10 µM MAN193, after co-expression of mCherry-talin-1 and PDE4B. g) Talin-1-positive focal adhesions at the basal membrane visualised with mCherry-talin-1 (561 nm excitation, 610-650 nm detection window). h) Merge of all channels. Scale bars, 20 µm.

Nonetheless, higher-resolution imaging revealed the emergence of distinct intracellular features superimposed on an otherwise diffuse cytosolic distribution (**Figure 3b**). Following analysis across multiple cells, two recurrent localisation patterns were consistently observed. First, co-labelling with the mitochondrial marker MitoTracker Deep Red revealed substantial mitochondrial localisation, confirming that MAN194 has a significant degree of colocalisation with the mitochondrial network (**Figure 3c-e). Figure 3c** also shows an overall increased signal within the nucleus of the cell (marked by Hoechst in **Figure 3d**), suggesting accumulation of the probe within intracellular sequestration compartments. Second, confocal imaging near the basal plasma membrane of PDE4B-overexpressing cells revealed localised accumulation of MAN194 at discrete peripheral structures (**Figure 3f**). Co-expression of mCherry-talin-1 identified these structures as talin-1-positive focal adhesions (**Figure 3g-h**).

To determine whether the observed localisation patterns reflected specific PDE4 binding, we preincubated cells with a saturating concentration of rolipram prior to MAN193 loading. Under these conditions, PDE4 binding sites are occupied by unlabelled rolipram, thereby preventing intracellular binding of MAN194 and allowing discrimination between specific and non-specific localisation patterns. Co-labelling with MitoTracker (**Figure 4b**) demonstrated persistent mitochondrial accumulation of MAN194 (**Figure 4c**), indicating that this localisation is largely independent of PDE4 binding. In contrast, rolipram preincubation abolished MAN194 accumulation at talin-1-positive focal adhesions (**Figure 4d-f**), demonstrating that the MAN194 signal observed in **Figure 3f** ireflects a specific population of PDE4 localised at these sites. This observation is consistent with previous reports describing recruitment of PDE4D5 to integrin adhesion complexes (Yarwood, Parnell, and Bird 2017; Yun et al. 2016), as well as the role of talin as an A-kinase Anchoring protein (AKAP), that scaffolds protein kinase A signalling (Kang et al. 2024).

**Figure 4.**
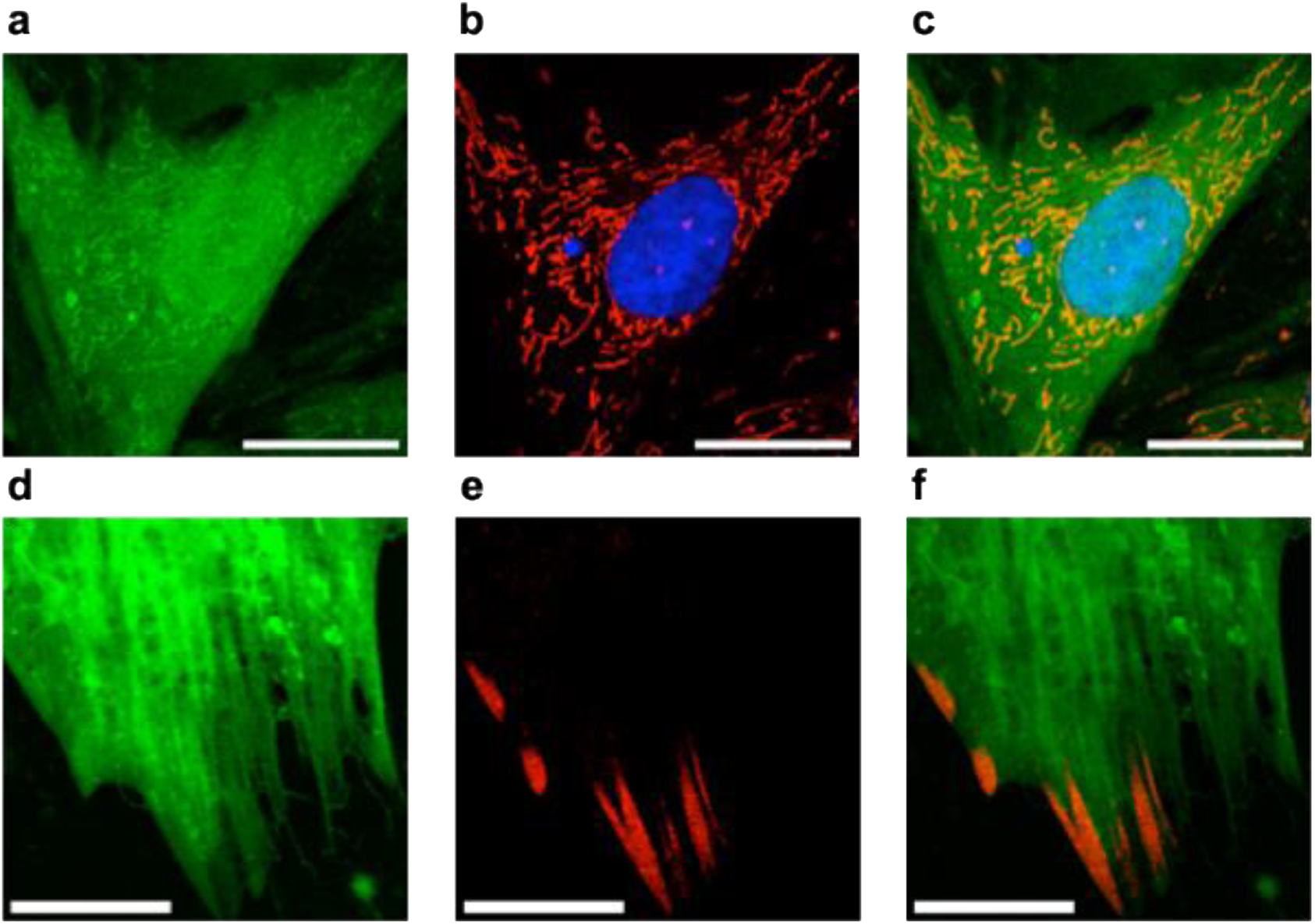
Live-cell uptake of MAN193 and localisation of MAN194 in H9c2 cells following rolipram pretreatment. a) H9c2 cells preincubated with 100 µM rolipram and imaged following 30 min incubation with 10 µM MAN193, further incubated with b) MitoTracker Deep Red (633 nm excitation, 640-700 nm detection window) and Hoechst (405 nm excitation, 430-470 nm detection window). c) Merge of the three colours. d) Confocal fluorescence images of H9c2 cells expressing mCherry-talin1 and PDE4B. Cells were preincubated with 100 µM rolipram and then loaded with 10 µM MAN193 for 30 minutes before imaging (488 nm excitation, 500-560 nm detection window). e) mCherry-Talin1 images from the same cells (561 nm excitation, 610-650 nm detection window). f) Merge of both channels. Scale bars, 20 µm.

Additional evidence that mitochondrial accumulation is largely non-specific was obtained by loading cells with FDA alone. Cells treated with 1 µM FDA displayed pronounced mitochondrial localisation (**Supplementary Figure 3a-d**), which remained detectable at concentrations as low as 100 nM (**Supplementary Figure 3e-h**). Because fluorescein is not expected to readily cross phospholipid bilayers (Loewenstein and Kanno 1964), including mitochondrial membranes, the mitochondrial signal observed following MAN193 loading is likely to arise from non-hydrolysed precursor molecules. One possible explanation is that high intracellular concentrations of MAN193 transiently saturate cytosolic esterase activity, thereby allowing uncleaved probe to access mitochondria before undergoing hydrolysis by mitochondrial esterases within the intermembrane space or matrix.

**Supplementary Figure 3.**
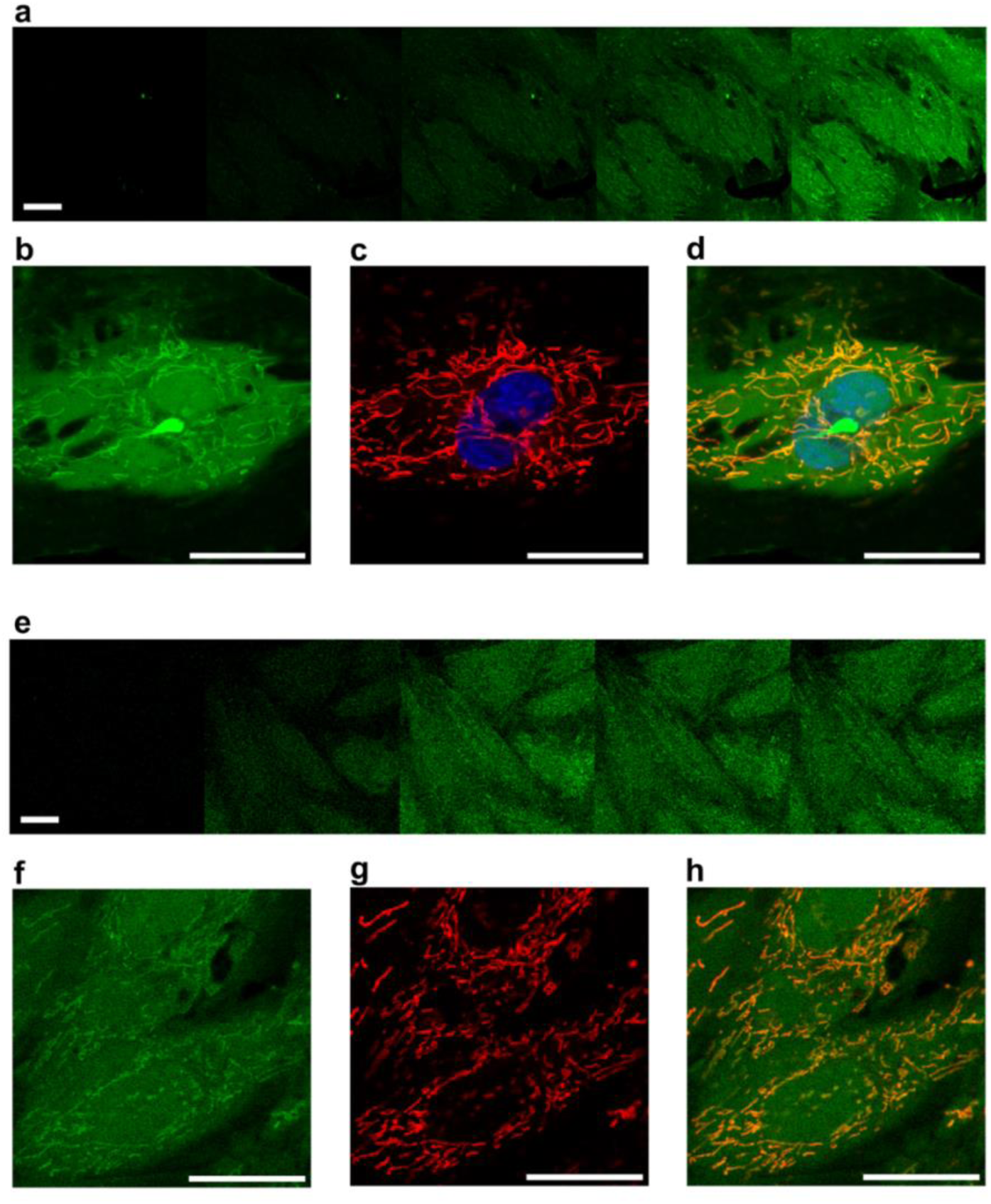
Uptake of fluorescein diacetate (FDA) in H9c2 cells. a) Time course of 1 µM FDA addition and intracellular conversion into fluorescein within H9c2 cells, monitored by confocal microscopy (100 Hz linescan rate, 488 nm excitation, 500-600 nm detection window). Each frame is acquired every approximately 410 s. b) Zoom-in cells imaged after 30 min incubation with 1 µM FDA, further incubated with c) MitoTracker Deep Red (633 nm excitation, 640-700 nm detection) and Hoechst (405 nm excitation, 430-470 nm detection window). d) merge of images in b) and c). e) Time course of 100 nM FDA addition and uptake in H9c2 cells. The representative mosaic shows confocal micrographs that are acquired every approximately 410 s. f) Cells imaged after 30 min incubation with 100 nM FDA, further incubated with g) Mitotracker Deep Red (633 nm excitation, 640-700 nm detection). h) combination of the images. Scale bars, 20 µm.

Based on these observations, hoping to mitigate this effect, we generated a different fluorescent analogue of rolipram (AG3481, rolipram-JF646, **Appendix 1**), employing the fluorogenic and cell membrane permeable dye JF646, originally derived from Silicon Rhodamine (Zheng et al. 2019). AG3481 was prepared using a synthetic approach analogous to that employed for MAN193 (see **Appendix 1)**. Interestingly, imaging in cells preincubated with saturating concentrations of rolipram confirmed that AG3481 displayed a marked non-specific localisation to mitochondria as well to perinuclear regions, that can be associated to the endoplasmic reticulum (**Supplementary Figure 4a**), albeit not to the nucleus. JF646 alone displays a localisation to the mitochondrial network and the nucleus (**Supplementary Figure 4b**). The additional non-specific localisation of AG3481 to the endoplasmic reticulum is evident when comparing uptake of rolipram-JF646 to MAN193 in cells preincubated with saturating concentration of rolipram, where AG3481 displays a perinuclear localisation that MAN194 does not show, whereas they both display mitochondrial localisation (**Supplementary Figure 4c-e**).

**Supplementary Figure 4.**
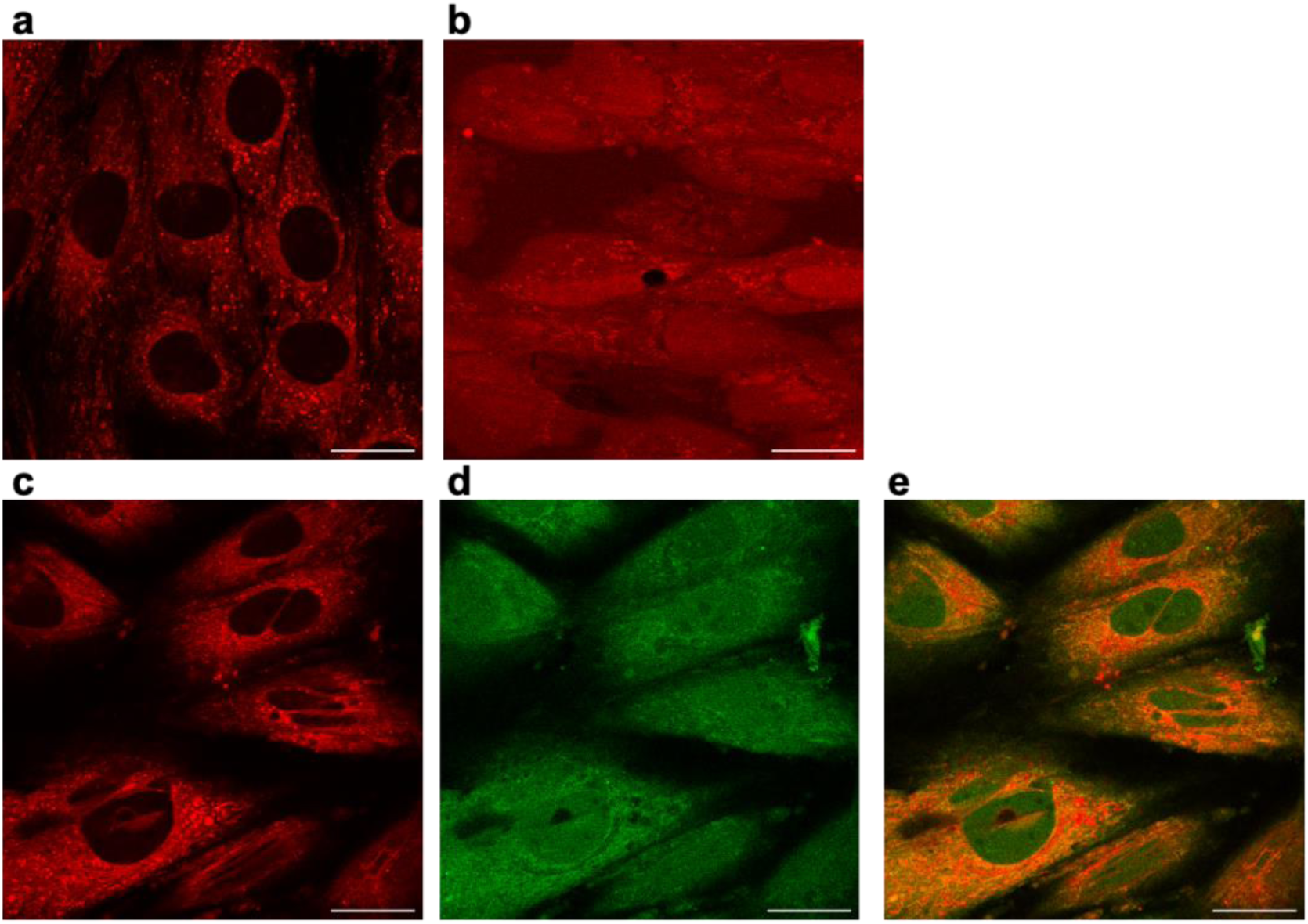
Uptake and localisation of AG3481 and JF646 in H9c2 cells. a) H9c2 cells preincubated with 100 µM rolipram prior to loading with 10 µM AG3481. Cells were imaged 13 minutes after probe addition. b) H9c2 cells loaded with 1 µM JF646. Cells were imaged 13 minutes after loading. Imaging was conducted at 100 Hz linescan rate, 633 nm excitation, 640-700 nm detection window on a Leica SP8 confocal microscope. c-d) H9c2 cells preincubated with 100 µM rolipram and then loaded with c) 1 µM AG3481 or d) 1 µM MAN193. e) Merge of the colour channels.

Since we did not observe significant advantages from using a next generation dye such as JF646 compared to FDA, we decided to further investigate our ability to use MAN193 as a reporter of intracellular rolipram binding. While overall intracellular abundance levels of PDE4 are not reported, assuming copy-number expression of all PDE4 isoforms in the 10^5^-10^6^ molecules per cell, together with a typical cellular volume of ∼1 pL, correspond to effective intracellular concentrations of 0.1-1 µM. Loading the cells with 10 µM of MAN193 will therefore lead to a fraction of free intracellular MAN194, making it difficult to determine its intracellular binding in the cytosol (**Figure 3c,f**). Loading cells with lower concentrations of MAN193, for instance 1 µM (**Supplementary Figure 5a-d**) or even 100 nM (**Supplementary Figure 5e-h**) appears to mitigate the issue of mitochondrial localisation.

**Supplementary Figure 5.**
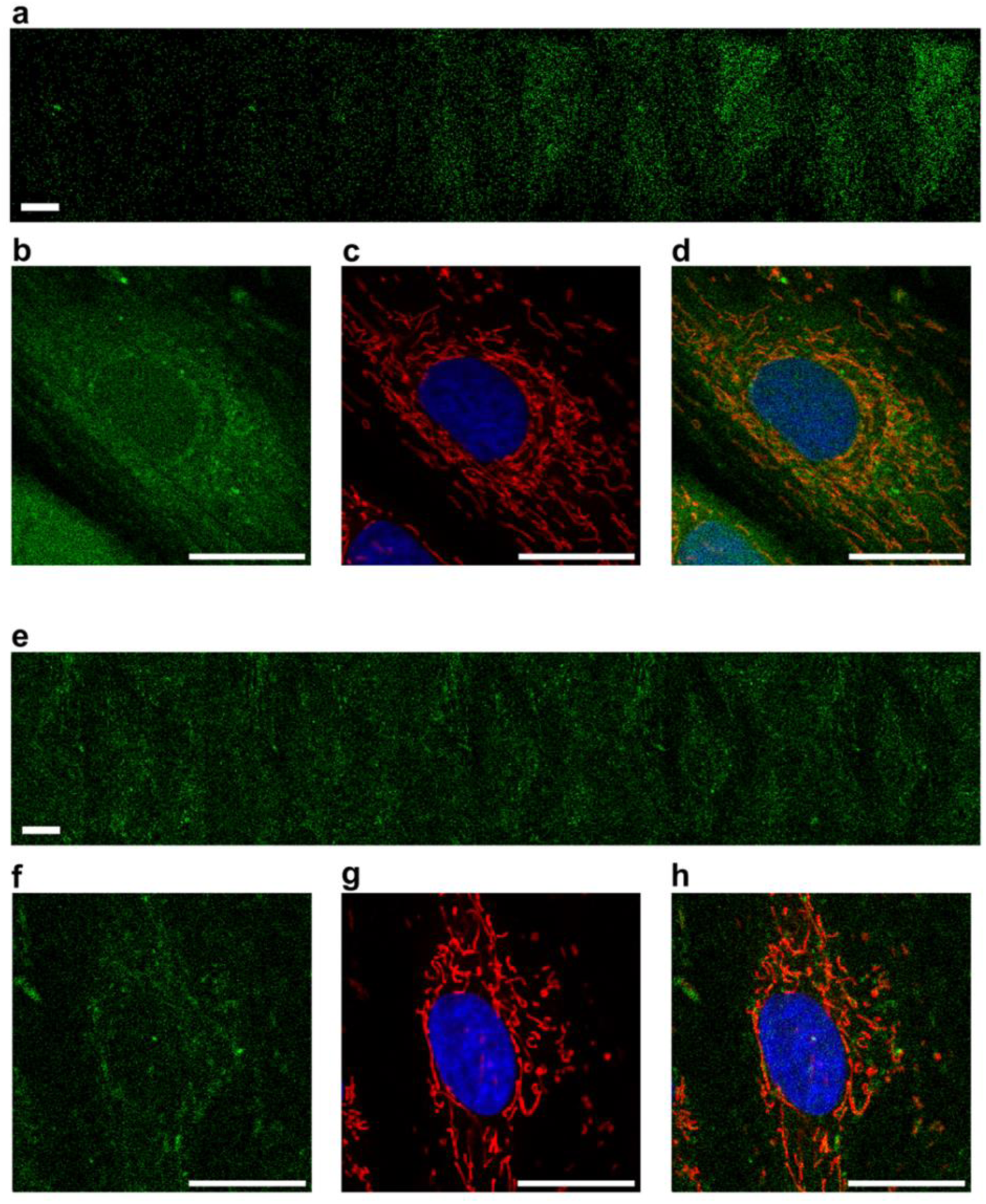
Concentration-dependent uptake of of MAN193 in live H9c2 cells using confocal microscopy. a) Time course upon addition of 1 µM MAN193 and intracellular conversion into MAN194 within H9c2 cells, monitored using confocal microscopy (100 Hz linescan rate, 100 nm pixel size, 488 nm excitation, 500-600 nm detection window). Frames shown in the mosaic are acquired every 410 s. b) Zoom-in cells imaged after 30 min incubation with 1 µM MAN193, further incubated with c) MitoTracker Deep Red (633 nm excitation, 640-700 nm detection) and Hoechst (405 nm excitation, 430-470 nm detection window). (d) merge of images in b) and c). e) Time course of 100 nM MAN193 addition and uptake in H9c2 cells. The representative mosaic shown is obtained by averaging every 10 frames and then taking every second frame from the new series. f) Cells imaged after 30 min incubation with 100 nM MAN193, further incubated with g) MitoTracker Deep Red (633 nm excitation, 640-700 nm detection) and Hoechst (405 nm excitation, 430-470 nm detection window). h) combination of the images. Scale bar, 10 µm. The time scale is indicated in the respective panels.

However, reducing probe concentration also leads to a substantial degradation in signal-to-noise ratio (**Supplementary Figure 5b,f**), prompting us to employ more selective fluorescence spectroscopy approaches to monitor intracellular binding and subcellular distribution beyond intensity-based imaging alone.

To monitor intracellular binding of MAN194, we therefore employed here a two-pronged approach, based upon i) image fluctuation spectroscopy to assess cytosolic binding of MAN194 and (ii) (epi-)fluorescence anisotropy imaging to monitor overall cellular binding. Beacuse fluorescence intensity alone cannot distinguish between free and PDE4-bound MAN194 we reasoned that intracellular binding could instead be inferred from changes in the spatial distribution of the fluorescent analogue. Previous studies have reported localised subcellular distribution of PDE4 isoforms (Jin et al. 1998) (Müller, Engels, and Fozard 1996) (Houslay and Adams 2003) (Taskén et al. 2001) (Fertig and Baillie 2018) (Pryzwansky, Kidao, and Merricks 1998) (Wills et al. 2016), suggesting that multiple PDEs may be concentrated within a diffraction-limited intracellular domains rather than uniformly distributed throughout the cytosol. In addition, our previous work identified a dimeric fingerprint for PDE4A1 (Bock et al. 2020), leading us to anticipate that leading us to anticipate that MAN194 would transition from a relatively monodisperse cytosolic population to a more spatially clustered distribution upon PDE4 binding. Such changes can be detected using confocal image fluctuation spectroscopy approaches, including molecular brightness analysis (Işbilir et al. 2021), (Digman et al. 2008), which quantify fluctuations in the variance of pixel intensity distribution in a region, or in a pixel over time (**Figure 5a**).

**Figure 5.**
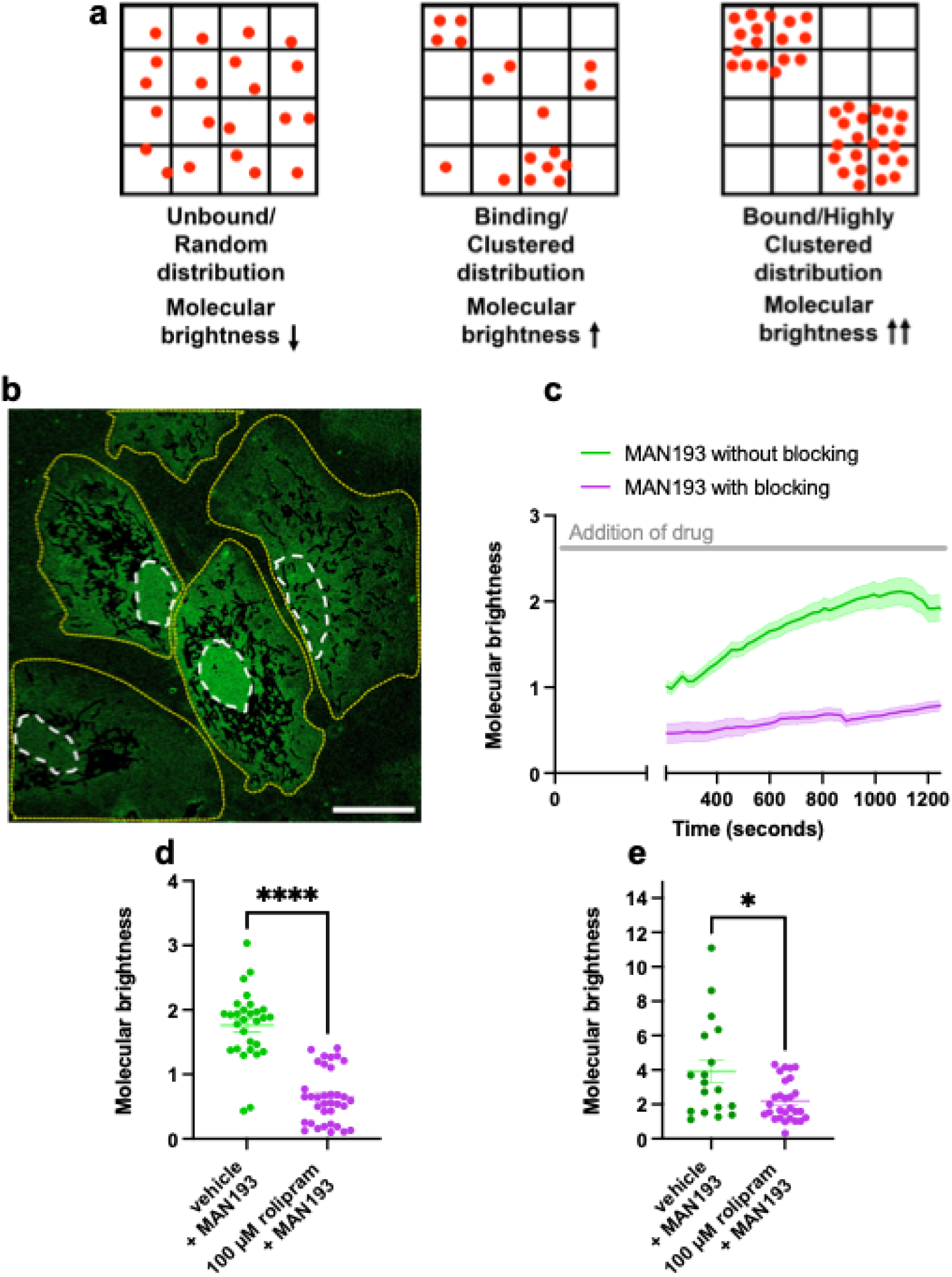
Molecular brightness analysis reveals the intracellular binding of MAN194 in H9c2 cells. a) Schematic representation of the molecular brightness concept applied to monitor intracellular binding of the fluorescent analogue. b) Representative processed confocal micrograph image used for molecular brightness analysis. Regions of interest (ROIs) generated using Cellpose Trackmate are outlined in yellow, and the nucleus is outlined in white. c) Molecular brightness evolution over time measured upon loading MAN193 in H9c2 cells preincubated with vehicle (green, n = 29) compared to cells pre-treated with 100 µM rolipram (violet, n = 35). Data are mean ± s.e.m. from n =3 independent experiments. d) molecular brightness values at ∼11 minutes (683 s) after the addition of MAN193 to the H9c2 cells compared between untreated (n = 29) and pre-treated (n = 35) cells. Results represented as dot plots with mean ± s.e.m from n = 3 independent experiments. e) Molecular brightness analysis of H9c2 cells overexpressing PDE4B, selected using a red fluorescent mRuby expression reporter. Data measured 30 minutes after addition of MAN193, without blocking (n = 18) and with blocking (n = 28) conditions. Results shown as dot plots with mean ± s.e.m from n = 2 independent experiments. An unpaired t-test was performed to determine significance (* P ≤ 0.05, **** P ≤ 0.0001).

We conducted molecular brightness analysis in H9c2 cells loaded with 10 µM MAN193. The analysis was restricted to cytosolic regions outside mitochondria, identified by their elevated fluorescent intensity (**Figure 3c-e**), as well as regions adjacent to the plasma membrane, where focal adhesion-associated PDE4 recruitment and pronounced MAN194 clustering were observed (**Figure 3g-h**).

To exclude mitochondrial signal, cells were counterstained with MitoTracker Deep Red, and a segmentation mask was applied to remove the mitochondrial network from the analysis, (**Figure 5b** and **Supplementary Figure 6).** Molecular brightness within the cytosol increased progressively following addition of MAN193, consistent with intracellular binding of MAN194 to spatially heterogeneous PDE4 populations. In contrast, cells preincubated with 100 µM rolipram prior to MAN193 addition showed no detectable increase in molecular brightness over time, consistent with competitive inhibition of MAN194 binding to PDE4 (**Figure 5c**). This difference was further confirmed by comparison of molecular brightness values measured 5 minutes after probe addition (**Figure 5d**). A similar increase in molecular brightness was observed in cells selectively overexpressing PDE4B (**Figure 5e**), the isoform used in the *in vitro* binding assays shown in **Figure 1d**. Together, these data suggest that cytosolic PDE4 populations, including PDE4B, are organised within sub-diffraction intracellular clusters rather than uniformly distributed throughout the cytosol. MAN194 binding to these clusters causes an increase in molecular brightness due to higher heterogeneity of pixel-to-pixel fluorescent intensities, compared to the homogeneously diffuse MAN194 in the cytosol observed following rolipram pretreatment.

**Supplementary Figure 6.**
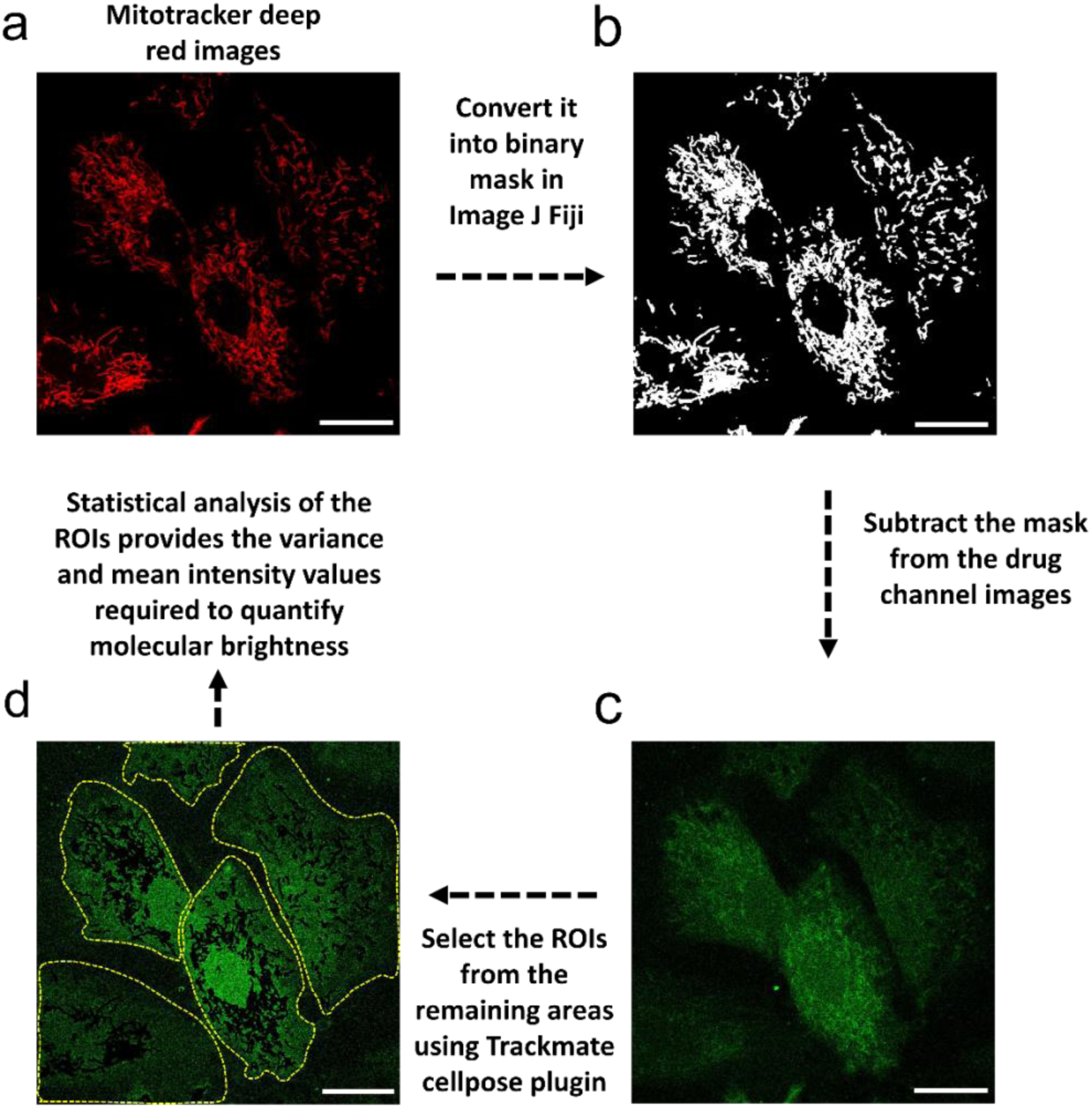
Workflow for mitochondrial signal removal prior to molecular brightness analysis. The MitoTracker Deep Red channel from the confocal micrograph a) was converted into a binary mask b). The binary mask was subtracted from the fluorescein channel of the micrograph c) and resulted in a new micrograph without the mitochondria. Cellpose TrackMate plugin in ImageJ Fiji was used to track cells across a kinetic series and ROIs were generated d). These ROIs and the new micrographs were fed into a custom-written ImageJ macro to obtain variance and mean values, which were used to quantify molecular brightness.

Similar results can be qualitatively obtained in HEK293 cells, albeit in these cells the cytosolic area visible in confocal cross sections without mitochondria is much smaller than in H9c2 cells (**Supplementary Figure 7**).

**Supplementary Figure 7.**
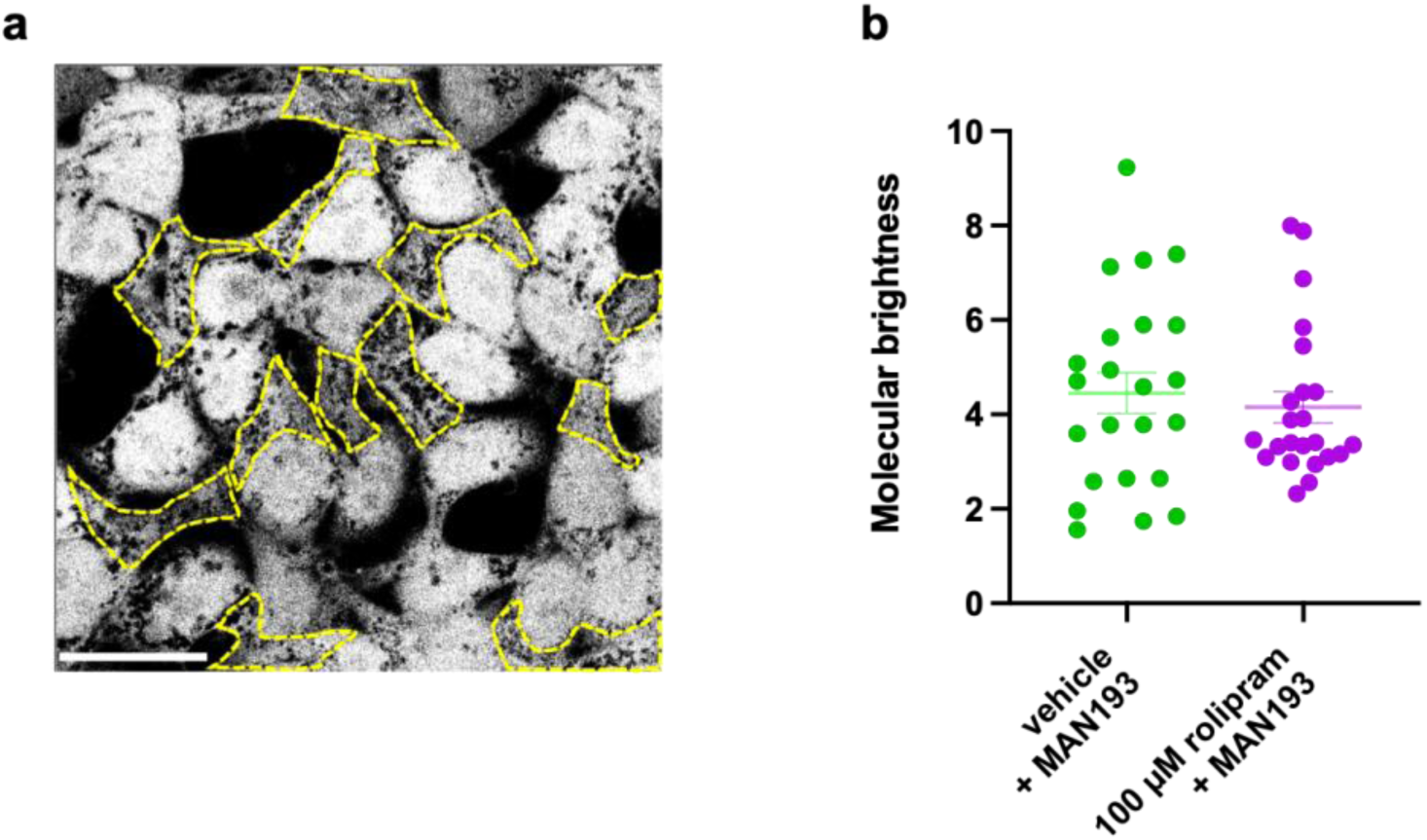
Molecular brightness analysis in HEK293T cells overexpressing PDE4B protein. HEK293 cells demonstrate significantly lesser cytoplasmic area and have a large nucleus, making it difficult to perform variance to mean calculations on an adequate number of pixels. (a) Selection of areas excluding the nucleus as ROIs. (b) Molecular brightness values calculated for MAN193 without blocking (n = 22) and MAN193 with blocking using 100 µM rolipram (n = 22). Result is represented as dot plot with mean ± s.e.m from n = 1 experiment.

While molecular brightness is a powerful tool, it is best suited to regions of the cell where the fluorescence signal is homogeneous, such as the cell cytosol (Annibale and Lohse 2020) and therefore requires exclusion of highly enriched MAN194 aggregates. We therefore complemented these measurements with fluorescence anisotropy imaging, which enables analysis across the entire cell. In this approach, samples were illuminated using linearly polarised excitation light under epifluorescence illumination, and fluorescence emission was recorded along two orthogonal polarisation directions. This approach (as discussed in **Methods**) allows obtaining a pixel-by-pixel average anisotropy value, and in turn reconstructing an anisotropy image (**Supplementary Figure 8**). Because fluorescence anisotropy values reflect the rotational diffusion of fluorescent molecules, increases in anisotropy indicate a larger fraction of probe molecules in bound states. Importantly, anisotropy measurements are independent of local fluorophore concentration and therefore largely decoupled from fluorescence intensity variations. We validated our imaging setup by comparing readouts from the microscopy images to those obtained using a fluorometer in cuvette. **Supplementary Figure 9a** shows a Perrin’s plot measured for fluorescein in the two setups. The results are further independent of the objective lens used (**Supplementary Figure 9b**) and the limiting anisotropies of MAN194 and fluorescein are comparable (**Supplementary Figure 9c**).

**Supplementary Figure 8.**
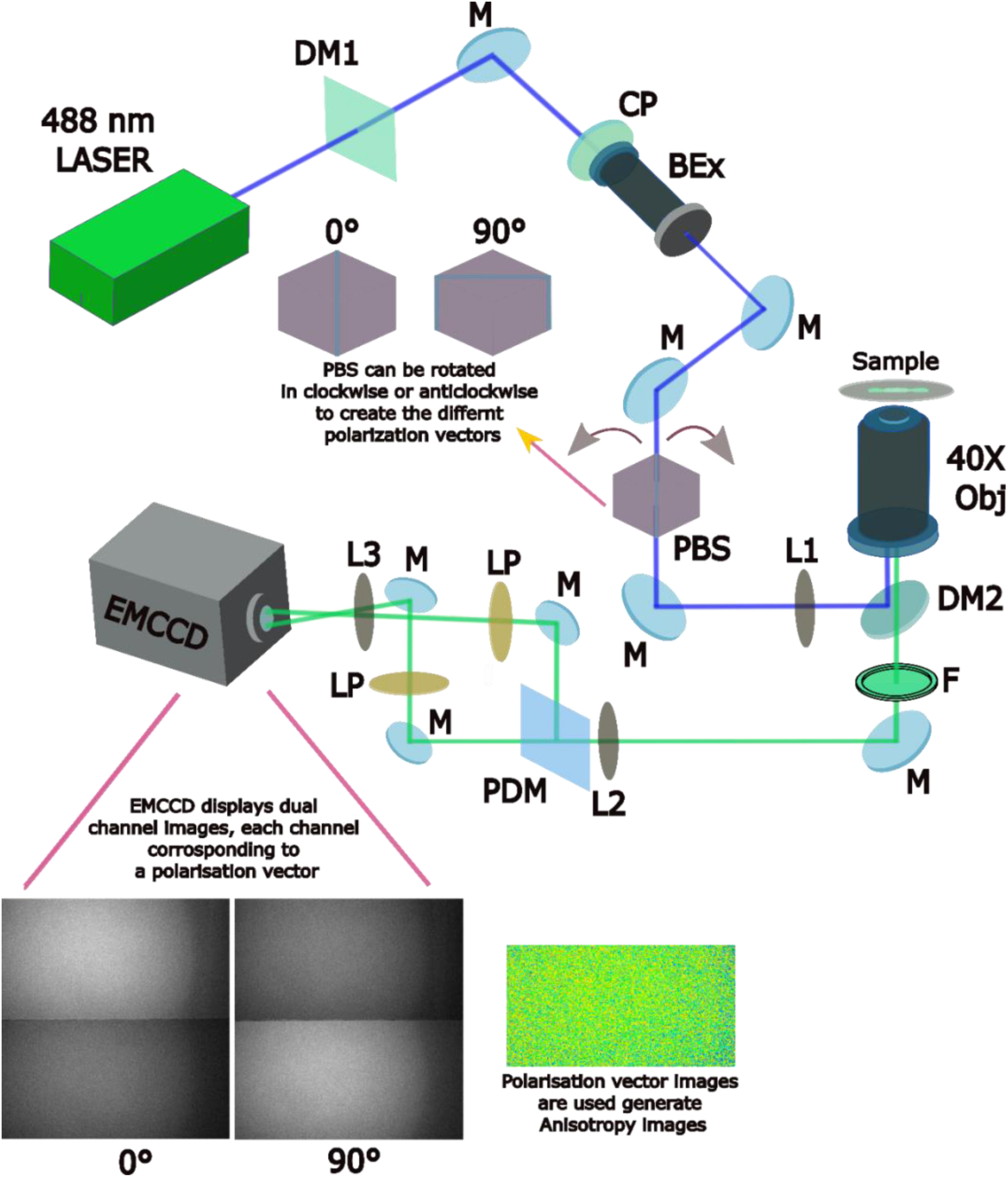
Microscopic setup used for fluorescence anisotropy imaging. Schematic representation of the custom-built microscope used for fluorescence anisotropy imaging. The scheme shows the excitation path (marked with the blue laser line) and emission path (marked with the green emission line). The excitation beam is passed through a circular polariser and expanded before passing through a polarising beam splitter (PBS) which converts the excitation beam into linearly polarised light which can be rotated clockwise or anticlockwise to obtain the two excitation polarisation vectors (parallel and perpendicular) required for calculation of anisotropy images. The excitation beam is then focused on the back focal plane of the objective lens using an achromatic doublet (L1, f=250) The excitation and emission from the sample are separated through a dichroic mirror (DM1, ZT 405/488/635) and the emission further selected through an emission filter (F, ET 572/26). The emission from the microscope is then steered to an Optosplit II (Cairn technology), where a polarising beamsplitter separates the emission into two separates spatially emission beams based on the polarisation direction of the emission. The separated polarised emission is further filtered using two linear polarisers and focused on two halves of the chip of the EMCCD camera. The camera thus displays dual channel images, each with its polarisation. The figure highlights the beam path as well as all the optical components: M = mirror, L = lens, DM = dichroic mirror, CP = circular polariser, Bex = beam expander, PBS = polarised beam splitter, Obj = Objective lens, F = emission filter, PDM = polarised dichroic mirror and LP = linear polariser.

**Supplementary Figure 9.**
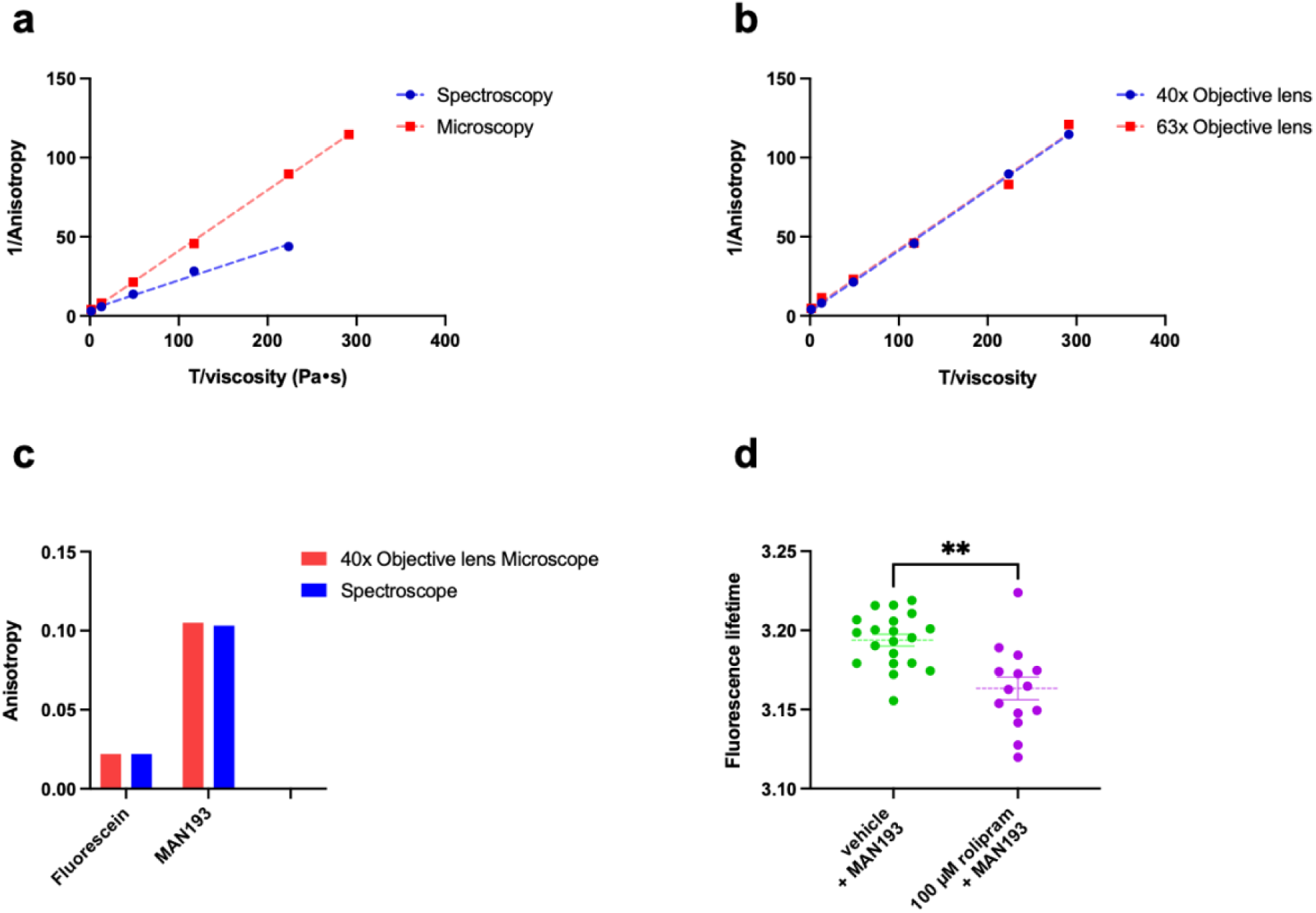
Fluorescence anisotropy and lifetime characterisation of the developed fluorescently labelled drug using microscopy. a) Instrumental validation of the developed fluorescence anisotropy imaging microscope by acquiring anisotropy readings using 100 nM fluorescein in 0-90% (w/w) glycerol solution and compared to values obtained from fluorescence spectrophotometer. Data is plotted as 1/anisotropy against T/viscosity and fitted using simple linear regression using GraphPad Prism. b) Perrin plots generated by using either the 40X or 63X objective lens. Data is plotted as 1/anisotropy against T/viscosity and is fitted using simple linear regression using GraphPad Prism. c) Limiting anisotropies (r_0_) calculated using 100 nM of Fluorescein or MAN194 measured using the developed anisotropy microscope and compared to fluorometer (spectroscope) measurements. Data is represented as bar chart and from n=1 experiment. d) Live cells fluorescence lifetime measurement following MAN193 loading in HEK293 cells without blocking (n = 14) or with blocking (n = 20). Results are represented as dot plots, along with mean ± s.e.m. and unpaired t-test was performed between the groups as statistical test (** P ≤ 0.01).

**Figure 6** shows three sets of fluorescence microscopy images, each comprising a transmitted light image (**Figure 6a-c**), a fluorescence intensity image **(Figure 6d-f**) and a false-colour fluorescence anisotropy image (**Figure 6g-i**). Individual anisotropy images for each sample analysed are shown in **Supplementary Figure 10**. Each set corresponds to HEK293T cells respectively pre-treated with vehicle (DMSO) and incubated for 30 minutes with 100 nM MAN193 (**Figure 6a,d,g**), pre-treated with 100 µM rolipram prior to incubation with 100 nM MAN193 (**Figure 6b,g,h**), or pretreated with vehicle and incubated with FDA (**Figure 6c,e,i**). Despite comparable fluorescence intensity distributions between vehicle- and rolipram-pretreated cells, anisotropy images, displayed in a false colour scale, revealed consistently higher anisotropy values in vehicle-treated samples, consistent with increased intracellular binding of MAN194.

**Figure 6.**
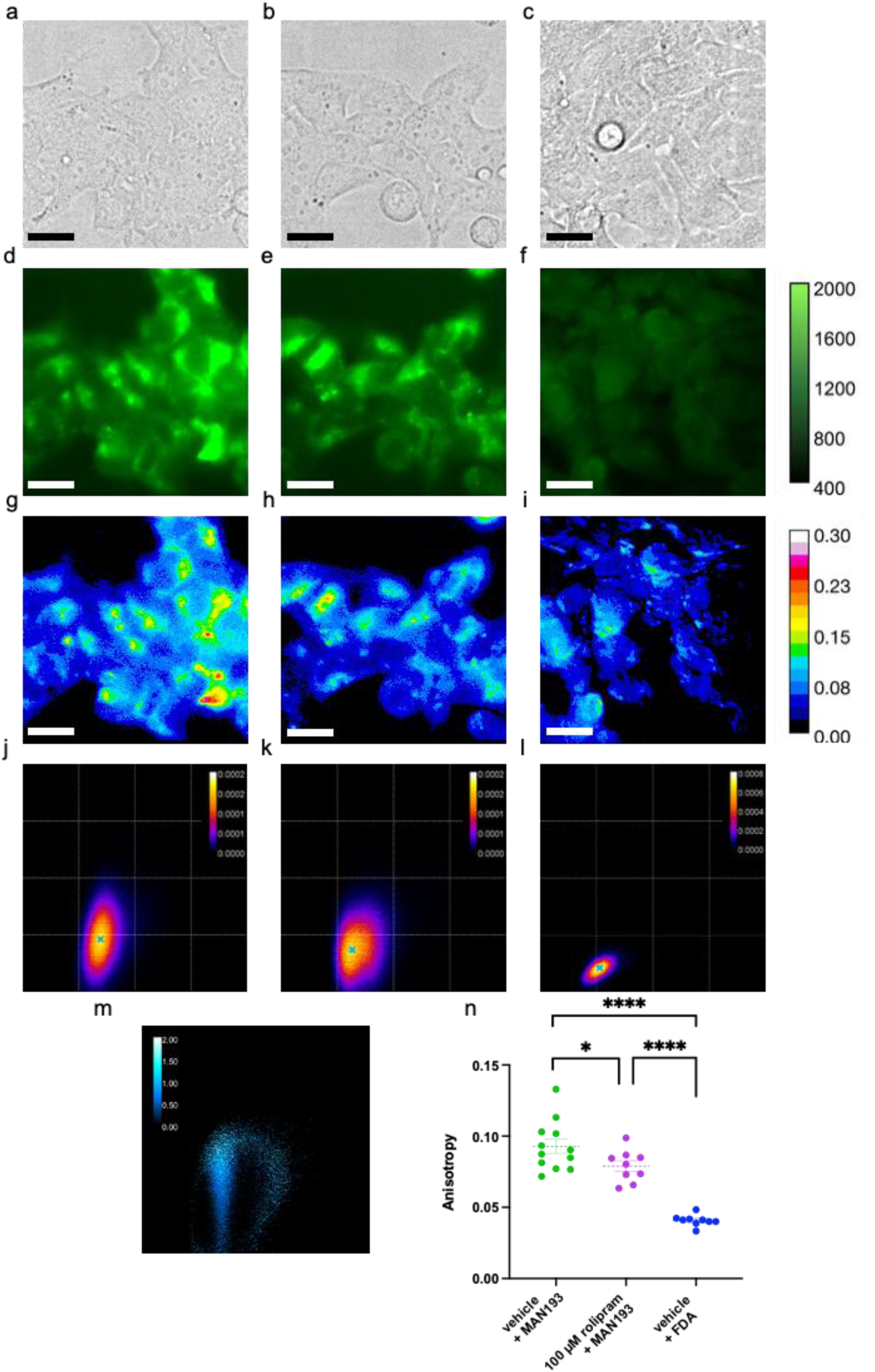
Visualisation of intracellular MAN194 binding using fluorescence anisotropy imaging. a-c) Representative brightfield images, d-f) fluorescence intensity images and g-i) false-colour fluorescence anisotropy mapsof HEK293 cells loaded with 100 nM MAN193 following pretreatment with either 1% DMSO vehicle control, 100 µM rolipram or fluorescein diacetate (FDA). The colour scale shows the related pixel intensity or anisotropy values in each condition. Scale bars, 20 µm. j-l) Averaged binned bivariate probability distribution plots generated from all experimental replicates across the three conditions. Each bin represents normalised pixels frequencies, providing a standard density distribution for comparison. The frequency of the pixel density is highlighted with the colour scale. The centroid of the distributions is marked with a blue cross, to highlight the highest probability value. m) Plot of the ratio between the number of normalised pixels in each bin for the averaged DMSO pre to RLP precondition. n) Comparison of the anisotropy values determined from the position of the peak in each of the binned bivariate probability distribution plots from the experimental replicates. The sample are either preincubated with 1% DMSO as vehicle control (n = 12) or 100 µM rolipram (n = 9) before MAN193 addition. Another set represents samples incubated directly with fluorescein diacetate (FDA, n = 9 coverslips; data is from n = 5 independent experimental replicates/days). Results are represented as dot plots, along with mean ± s.e.m. and unpaired t-test was performed between the groups as statistical test (* P ≤ 0.05, **** P ≤ 0.0001).

**Supplementary Figure 10.**
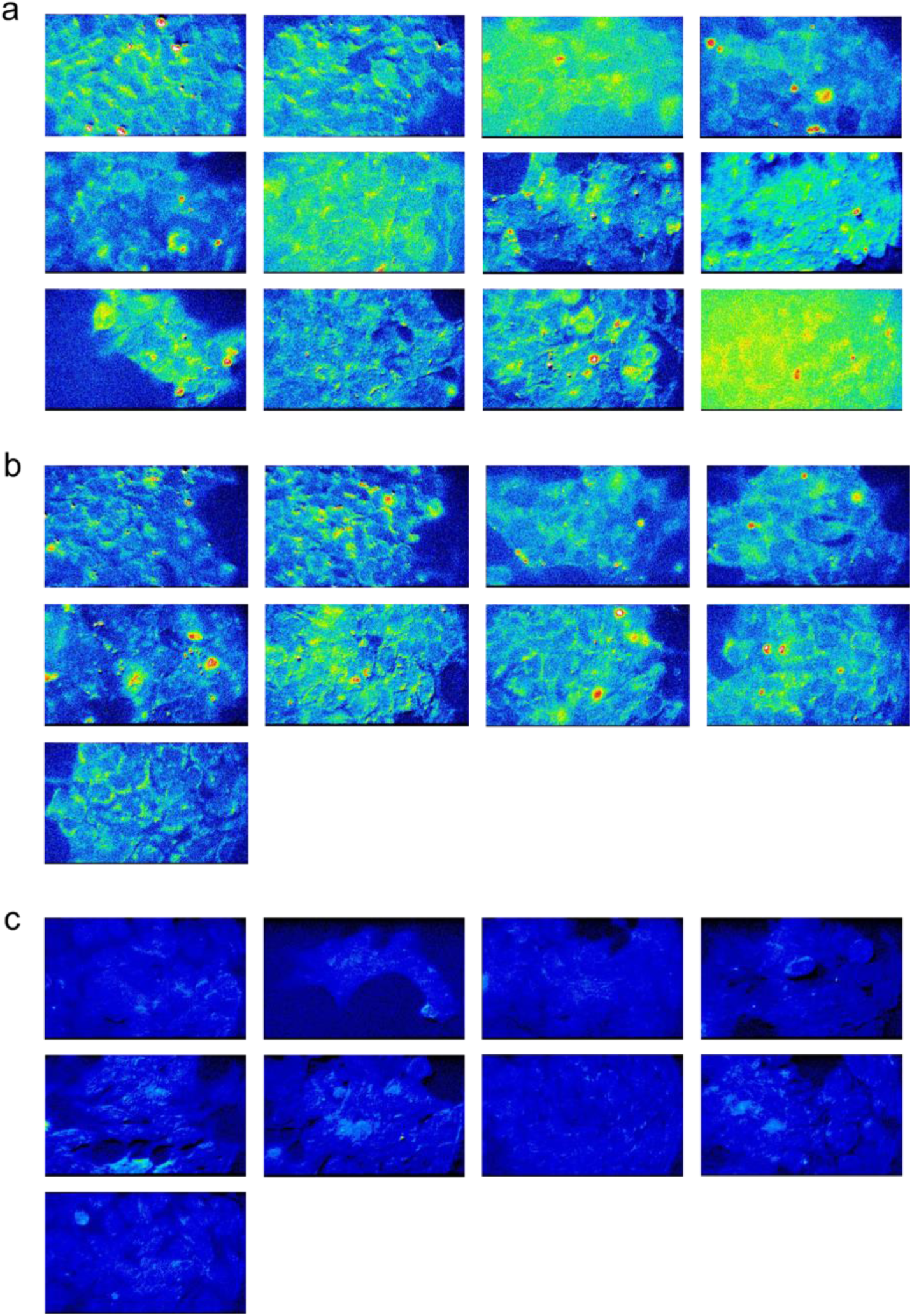
Fluorescence anisotropy imaging datasets. Anisotropy images across experimental conditions. Panels in a) and b) display the anisotropy images from samples incubated with MAN193 without blocking and MAN193 with blocking respectively. Panels in c) show the anisotropy images of samples incubated with FDA.

To quantitatively analyse these data, we conducted generating Binned Bivariate Probability Distribution plots, which are 2D histograms of the pixel anisotropy values (*r)* in each pixel *vs* the pixel intensity (*I)* (**Supplementary Figure 11)**. The overall 2D *r* vs *I* histograms highlight an increase of r values -for comparable *I* values-between the vehicle and rolipram pre-incubated controls. The combined histograms from multiple experiments are shown in (**Figure 6j-l**), and the 2D *r vs I* histograms from each sample are shown in **Supplementary Figure 12**. Fluorescein displays an overall much lower fluorescence anisotropy *r*.

**Supplementary Figure 11.**
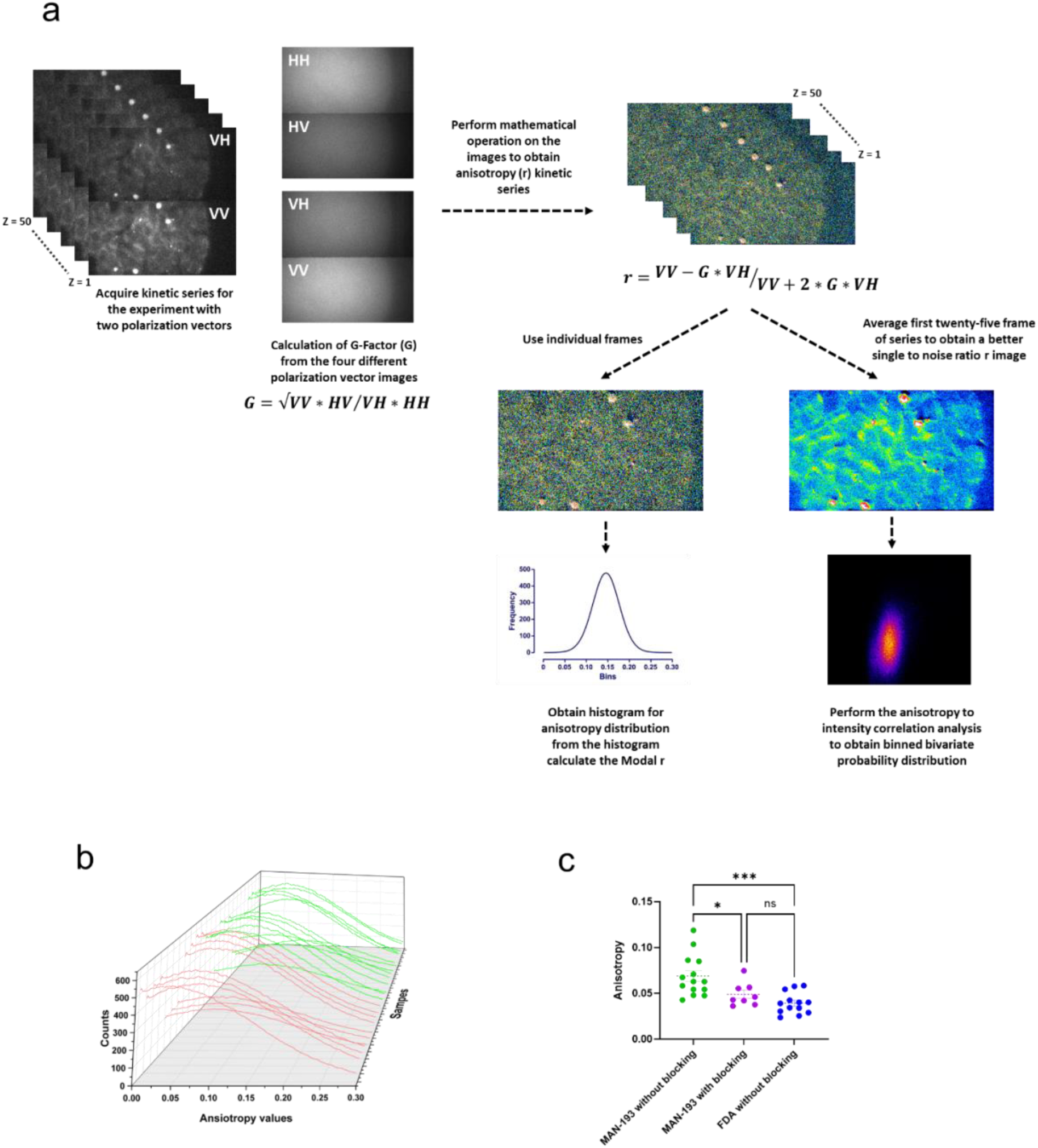
Fluorescence anisotropy imaging and analysis protocol. a) Schematic representation of the fluorescence image analysis protocol. A kinetic series for each sample is obtained and containing dual images, which represent VH and VV polarisation respectively. G-factor was calculated from four polarised images, HH, HV, VH and VV obtained with the imaging setting used for respective conditions and by rotating the polarised beam splitter clockwise or anticlockwise. The sample was 100 nM fluorescein in 50% W/W glycerol solution as the sample. G-factor was mathematically operated into the VH and VV images from kinetic series using Image J to obtain the anisotropy series. This kinetic series was subjected to two different kinds of analysis. In the first case, mathematical operations were performed on each frame to obtain the anisotropy histogram and calculate the modal anisotropy values. In the second case, first 25 frames were averaged together to obtain a new anisotropy image with better signal to noise ratio. This image was subjected to intensity vs anisotropy correlation analysis and to further obtain binned bivariant probability distribution plots. b) The plot shows the anisotropy histograms from MAN193 without blocking in green and histograms from MAN193 with RLP blocking in red. These histograms were used to recover modal anisotropy value. c) Comparison of the modal anisotropy values averaged from first 10 frames of each kinetic movies for an experiment preincubated with 1% DMSO as vehicle control (n = 12) or 100 µM rolipram (n = 9), or fluorescein diacetate (FDA, n = 9) (data is from five independent experimental replicates). Results are represented as dot plots, along with mean ± s.e.m. and unpaired t-test was performed between the groups as statistical test (ns: non-significant, * P ≤ 0.05, *** P ≤ 0.001).

**Supplementary Figure 12.**
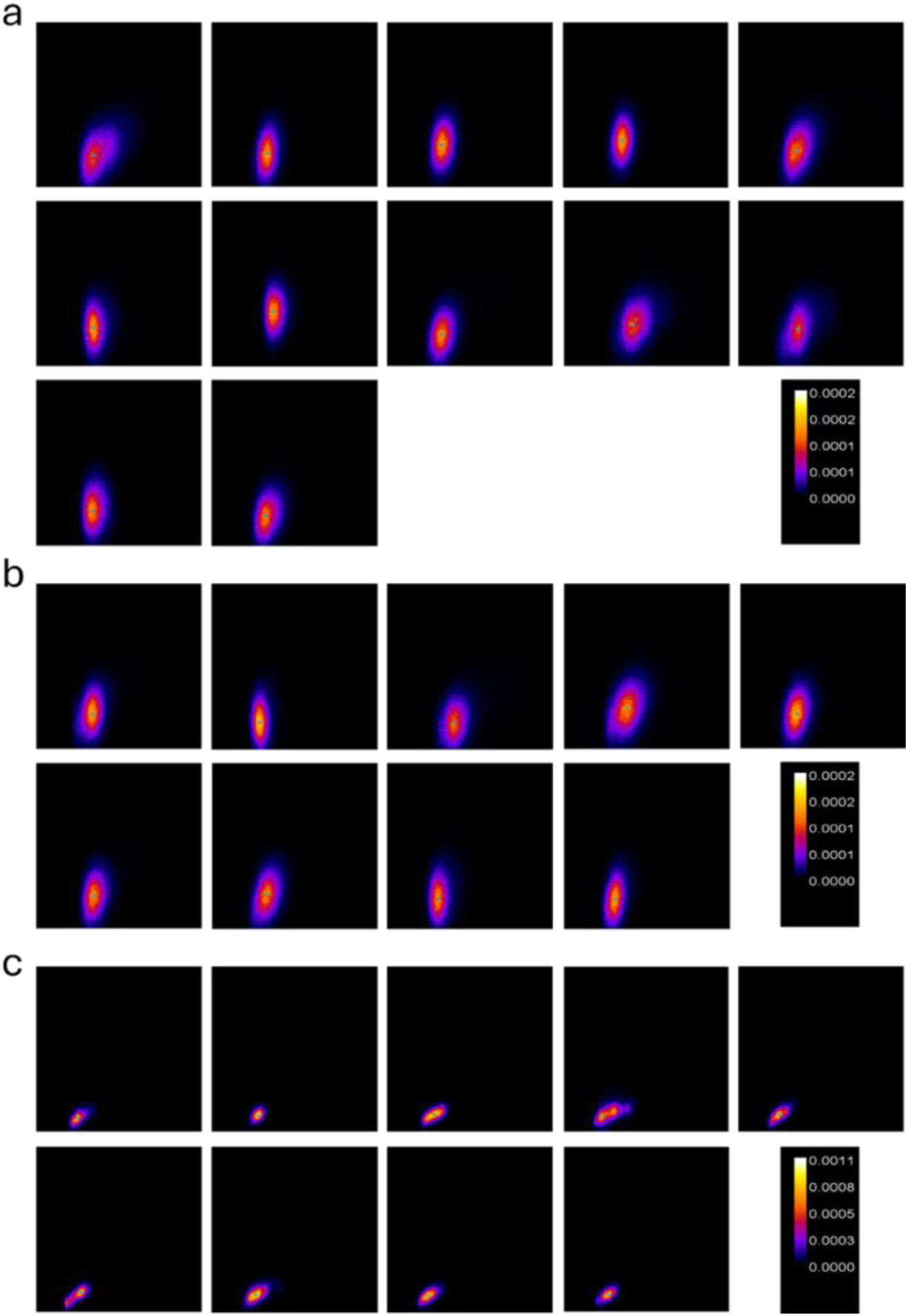
Binned bivariate probability distribution plots from each experiment. Representative 2D anisotropy vs intensity histograms containing the normalised distribution of pixels across experimental conditions. Panels in a) and b) display the plots from experiments where HEK193c cells were loaded MAN193 without vehicle preincubation or MAN193 after preincubation with 100 µM rolipram. Panels in c) show the plots from experiments from cells preincubated with FDA (without any pretreatment). The colour scale shows the frequency distribution of the pixels. The centroid of each distribution has been marked with a blue cross, indicating the maximum the probability distribution. The coordinates of the maxima were used to obtain the local anisotropy and intensity values for further analysis.

This difference is further illustrated by plotting the ratio of the overall 2D *r vs I* histograms obtained from vehicle-preincubated over rolipram-preincubated samples loaded with MAN193 (**Figure 6m**): it displays a plume increasing in strength for growing r values, at a defined intensity value, reflecting the fact that, on average, the anisotropy readouts of pixels of comparable intensity are higher in the vehicle samples (more MAN194 bound) as opposed to the rolipram-preincubated (less MAN194 bound). This is further quantitatively appraised by plotting the anisotropy value associated to the maximum of each 2D *r vs I* histogram **(Figure 6n)**, which illustrates how these values compare across the three datasets: significantly higher fluorescence anisotropy from the samples loaded with MAN193 after pre-incubation with vehicle than those pre-incubated with rolipram, or those samples loaded with FDA.

The ability to analyse fluorescence anisotropy as a function of fluorescence intensity (r versus I) provides a powerful spectroscopic tool for distinguishing simple intracellular accumulation of a fluorescent probe from regions associated with probe binding. In particular, this approach enables assessment of whether regions of elevated fluorescence intensity also display increased anisotropy consistent with restricted rotational diffusion and target engagement.

We used this analysis to determine whether the fluorescence accumulation of MAN194 observed at mitochondria reflected specific PDE4 binding or non-specific intracellular sequestration Representative fluorescence images of individual cells treated with vehicle + MAN193 (**Figure 7a**), rolipram + MAN193 (**Figure 7b**) or FDA alone (**Figure 7c**) all displayed pronounced a clear contrast and signal accumulation at mitochondria. Correspondinganisotropy maps likewise showed elevated anisotropy within mitochondrial regions across all conditions (**Figure 7d-f**).Analysis of anisotropy values across whole-cell regions of interest reproduced the population-level differences observed in **Figure 6**, with higher anisotropy values detected in vehicle-treated compared with rolipram-pretreated cells (**Figure 7g**). Nonetheless, it is now possible to conduct a cursor selection in the r vs I histogram from each cell’s image, and identify pixels belonging to specific features, in this case the mitochondrial network, and extract their anisotropy. **Figure 7h-j** illustrates the principle of cursor selection on a representative HEK293 cell, where pixels from two regions, cytosol and mitochondria, are masked by two different colours, respectively cyan and red (fluorescence intensity and anisotropy images from three cells from each condition are reported in **Supplementary Figure 13**).

**Figure 7.**
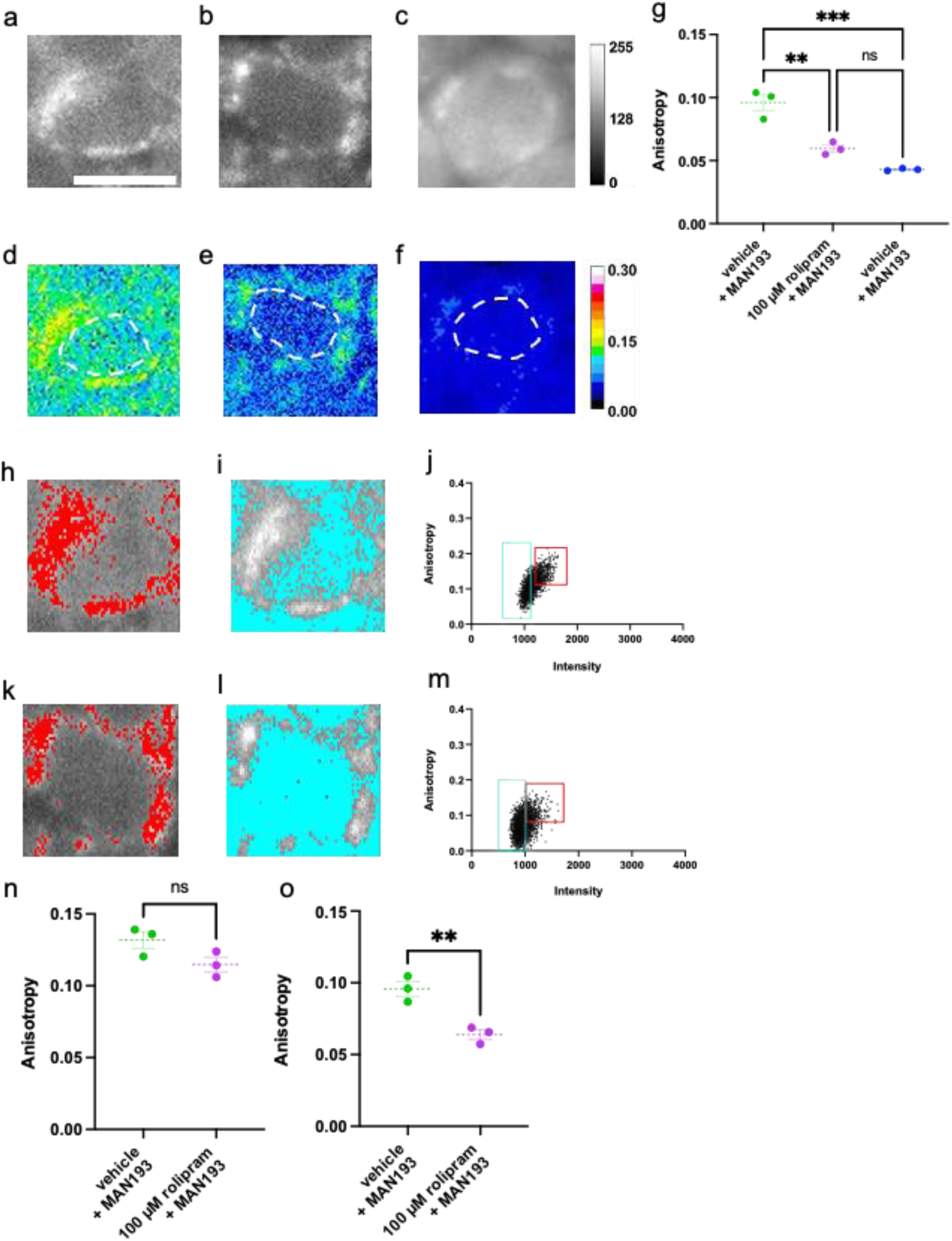
Subcellular fluorescence anisotropy analysis from single cell imaging. A-c) Representative fluorescence intensity image of a single cell loaded with MAN193 after preincubation with a) vehicle, b) 100 µM rolipram or c) FDA without any pretreatment. d,-f) Corresponding anisotropy images. The area marked with white dashed lines is used to calculate the mean anisotropy from each of the conditions. g) Mean anisotropy from each condition is represented as dot plots, along with mean ± s.e.m. An unpaired t-test was performed between the groups as statistical test (n = 3) (* P ≤ 0.05). h) Spatial selection of mitochondrial regions in the intensity image displayed in a (cell preincubated with vehicle), where the selected pixels are highlighted with a red mask. i) The non-mitochondrial cytosolic area is highlighted with cyan mask. j) The selections are based on the regions highlighted in the r vs I histograms. k) Spatial selection of mitochondrial regions in the intensity image displayed in b) (cell preincubated with 100 µM rolipram), where the selected pixels are highlighted with a red mask. l) The non-mitochondrial cytosolic area has been highlighted with cyan mask. m) The selections are based on the regions highlighted in the r vs I histograms. n) Comparison of anisotropy in the mitochondrial areas between cells preincubated with vehicle and cells preincubated with 100 µM rolipram. Anisotropy values are represented by dot plots with mean ± s.e.m. (n = 3). o) Comparison of anisotropy in the cytosolic areas between cells preincubated with vehicle and cells preincubated with 100 µM rolipram. Anisotropy values are represented by dot plots with mean ± s.e.m. (n = 3). Unpaired t-test was performed between the groups as statistical test (ns: not significant, ** P ≤ 0.01).

**Supplementary Figure 13.**
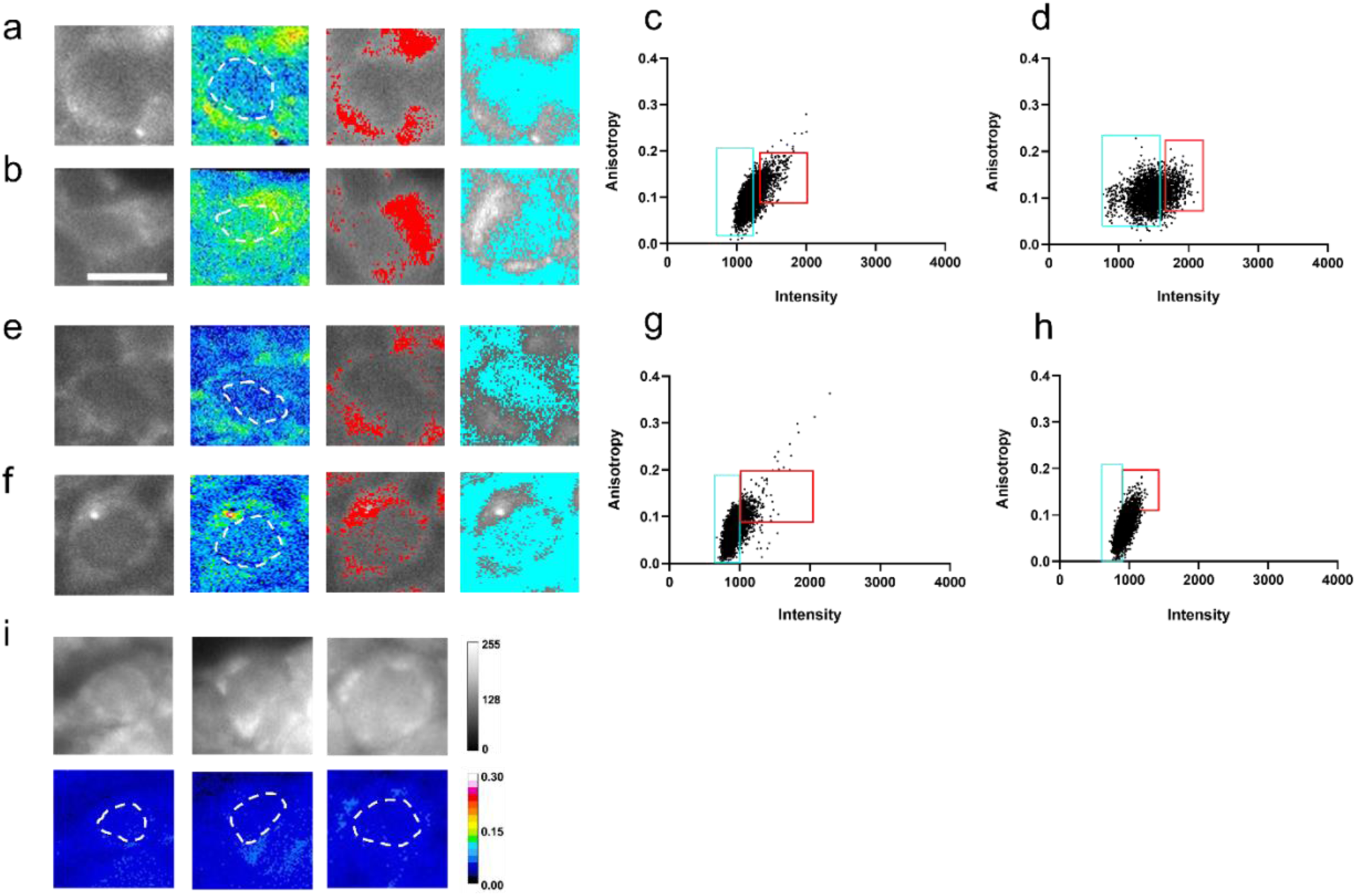
Subcellular localised binding of fluorescent drug in single cells. (a-b) Fluorescence intensity and anisotropy images of cells treated with MAN193 without blocking, with mitochondrial regions highlighted in red and non-mitochondrial regions in cyan. (c-d) corresponding anisotropy vs intensity correlation plots. (e-f) fluorescent intensity and anisotropy images along with the mitochondrial highlighted regions in red and non-mitochondrial regions highlighted in cyan associated to MAN193 with blocking along with the co-relation plot (g-h). (i) Fluorescent intensity images and corresponding anisotropy images from FDA without blocking. Colour scales indicate fluorescence intensity and anisotropy values. Scale bar, 20 µm.

This approach allows us to select pixels based on their intensity in the *r vs I* histogram, visually confirming that they match a feature in the image via the masking, and then extracting the local anisotropy values. We can then compare the mitochondrial and cytosolic fluorescence anisotropies from samples pretreated with vehicle (**Figure 7h-j**) or rolipram (**Figure 7k-m**). This allows us determining that a clear and significant anisotropy difference between the two conditions exists in the cytosol **(Figure 7n)**, in line with the results reported in **Figure 6**. On the other hand, no significant difference is present between the anisotropy readouts of MAN194 in the mitochondria **(Figure 7o)**, despite both fluorescence intensity and anisotropy being higher there. We interpret this result as restricted MAN194 rotational diffusion in the mitochondria, rather than specific PDE4 binding.

## Discussion

The ability to directly visualise intracellular drug–target engagement with subcellular resolution has long represented a major challenge in pharmacology and chemical biology. Here, we present a strategy to address this challenge through development of MAN193, a fluorogenic analogue of the PDE4 inhibitor rolipram, generated by conjugation to fluorescein diacetate (**Figure 1a**). This design exploits the fluorogenic character of FDA to create a cell-permeable, initially non-fluorescent prodrug that is converted intracellularly into the fluorescent form MAN194 by endogenous esterases, enabling selective visualisation of intracellular drug binding.

Biophysical characterisation demonstrated that MAN194 retains the core spectral properties of fluorescein (**Figure 1b,c**) while preserving specific binding to the PDE4 catalytic domain, albeit with reduced affinity (K_d_∼ 60 nM) compared to unlabelled rolipram (**Figure 1d**). Such affinity reductions are commonly observed following fluorophore conjugation of small-molecule ligands, including fluorescent GPCR ligands (Bresinsky et al. 2023; Bathe-Peters et al. 2021). Notably, in our previous work, conjugation of FDA to cAMP did not alter the *in vitro* dissociation constant of 8-F-cAMP from PKARI*α* (Bock et al. 2020), further reinforcing the rationale for using FDA as a fluorescent conjugate. Importantly, however, MAN193 retained cellular activity and behaved as a comparatively weak PDE4 inhibitor, a property that may prove advantageous for imaging applications by minimising perturbation of endogenous signalling dynamics.

A major advance of this study is the demonstration that fluorescent drug analogues can be used to visualise endogenous intracellular target populations in living cells without genetic manipulation or antibody delivery. Using MAN193, we identified a population of PDE4 localised at talin-1–positive focal adhesion complexes in cardiomyocyte-like cells. Although PDE4 recruitment to integrin adhesion complexes has previously been inferred biochemically and through fixed-cell imaging approaches(Serrels et al. 2010), our results demonstrate direct live-cell visualisation of endogenous PDE4-associated signalling domains.

This application is not readily accessible using classical immunofluorescence in living cells: commercially available anti-PDE4 antibodies have been validated for immunofluorescence imaging of fixed, permeabilised cells, enabling subcellular localisation studies of PDE4A, PDE4B and PDE4D isoforms (Jin et al. 1998; Taskén et al. 2001; Terrin et al. 2012). However, conventional IgG antibodies (∼150 kDa) are impermeant to intact plasma membranes, precluding their application in live-cell imaging of this intracellular target. Emerging nanobody-based intrabodies offer potential solutions but remain unavailable for PDE4 targets and require genetic manipulation of the cell (Traenkle et al. 2015). MAN193 therefore addresses a critical unmet need for dynamic, real-time imaging of PDE4 in living cells.

A central technical challenge in intracellular fluorescent pharmacology is distinguishing free intracellular probe from target-bound populations. We addressed this limitation using two complementary fluorescence spectroscopic approaches. Molecular brightness analysis exploits the tendency of PDE4 isoforms to form sub-diffraction-limit clusters in the cytosol (Bock et al. 2020). Binding of MAN194 to these clusters generates local intensity heterogeneities that are detected as an increase in molecular brightness compared to cells pretreated with the dark inhibitor rolipram (**Figure 5**). In parallel, fluorescence anisotropy imaging, provides a concentration-independent readout of intracellular binding based on restricted rotational diffusion. Using this approach, we detected rolipram-displaceable binding of MAN194 in the cytosol of living cells, providing strong evidence that the observed signal reflects specific PDE4 engagement (**Figure 6**).

An important finding of this study is the ability to distinguish between specific and non-specific accumulation of MAN194, and in general of fluorescent drug analogues. This was enabled by the combination of *r* vs *I* histogram analysis and spatial masking, highlighting the ability of anisotropy imaging to report not only localisation but target engagement with subcellular resolution (**Figure 7**). This distinction proved particularly important for interpreting mitochondrial accumulation of MAN194. Although both fluorescence intensity and anisotropy were elevated within mitochondria, anisotropy values were insensitive to rolipram competition, indicating that mitochondrial enrichment reflects restricted rotational diffusion and intracellular sequestration rather than specific PDE4 binding. More broadly, these findings highlight the importance of directly measuring intracellular target engagement rather than inferring binding solely from subcellular accumulation patterns.

While the accumulation of MAN194 to the mitochondria appears an undesired characteristic of our compound, most likely driven by FDA, it highlights the importance of monitoring the subcellular localisation and uptake of any new drug (Greenwood et al. 2019). Fluorescent drug analogues are well suited to address this challenge and could be incorporated as part of routine drug development pipelines.

An important implication of this work is that FDA despite its simplicity and long-standing use as a viability probe, performs comparably or better than newer fluorogenic scaffolds for intracellular drug conjugation. In contrast to the JF646-based probe, which displayed substantial additional endoplasmic reticulum accumulation, MAN193 retained a comparatively more restricted non-specific localisation. These results suggest that FDA-based fluorogenic conjugation may provide a practical and inexpensive strategy for generating fluorescent analogues targeting intracellular proteins.

Together, our findings establish fluorogenic drug conjugation combined with quantitative fluorescence spectroscopy imaging as a broadly applicable framework for mapping intracellular drug–target engagement with subcellular resolution. Extension of this strategy to additional pharmacophore classes and fluorophore scaffolds may enable systematic interrogation of the spatial pharmacology of small molecules in living systems.

## Acknowledgments

We are grateful to Elisabetta Armani (Chiesi Farmaceutici) for sharing the PDE4B catalytic domain protein. S.B. is grateful to Richard Edel and Remi Fritzen (University of St Andrews) for help and guidance related to molecular cloning.

## Author contributions

P.A. designed research

S.B., M.A.N., A.G., J.S., P.A. performed research

S.B., P.A., J.S., H.S. data analysis

B.T. G. shared reagents and research tools

S.B. graphical analysis and visualisation

P.A. and S.B. wrote the manuscript

A.G. and B.T.G contributed to writing the manuscript

P.A., A.G., H.S. supervision of experiments

P.A., A.G., P. G. C., H.S. funding acquisition

## Competing Interests

The authors declare no competing interests

**Data Availability Statement**

**Code Availability Statement**

## Materials and Methods

### Chemical synthesis

Details of the chemical synthesis and characterisation of the fluorescent analogues is described in Appendix A.

### Cell culture

The cell lines used for the study were HEK293T cells (ECACC 96121229, Sigma-Aldrich) and H9c2 cells H9c2 (ATCC; CRL-1446). HEK293T cells are immortalised human embryonic kidney cells widely used because of their ease of culture and transfection.” H9c2 cells are rat cardiomyoblast-derived cells (Hescheler et al. 1991). These cell lines were selected based on reported expression of PDE4 isoforms.

The cells were maintained in a T-25 or T-75 flasks in presence of DMEM high glucose media (Thermo Fisher Scientific) supplemented with 10% FBS (Thermo Fisher Scientific) and 1% P/S antibiotics (Thermo Fisher Scientific) (this solution will be referred to as cell-culture media henceforth) in a CO2 incubator at 5% CO2 and 37°C. Once the cells reached 80–90% confluency, cells were washed with 5 mL phosphate-buffered saline without Ca2+ and Mg2+ ions (Thermo Fisher Scientific) and trypsinised using 3 mL 0.05% trypsin/ ethylenediaminetetraacetic acid solution. Cells were passaged after every 2 to 3 days. Cells were routinely tested for mycoplasma infection using Venor®GeM OneStep kit (Minerva biolabs).

### UV-VIS spectrometry and fluorometry

Absorbance measurements of the compounds were performed using Evolution 350 UV-Vis Spectrophotometer (ThermoFisher scientific). The compounds were dissolved in TRIS buffer to a final concentration of 1 µM and a high precision cell quartz cuvette (Hellma Analytics) of path length 10 mm were used to record the measurements. Initially, the absorbance of TRIS buffer was measured and then the absorbance of molecules was obtained after subtracting the background data.

Excitation and emission spectra were recorded using the Cary 300 Fluorometer. The stocks of Fluorescein, MAN194 and AG3461 in were dissolved in a buffer made of 25 mM Tris-HCl (pH 8.0); 200 mM NaCl; 5 mM DTT, 0.5 mM EDTA. A final working concentration of 10 nM was used for measuring the excitation and emission spectra of the compounds. The emission spectra of the MAN194 and Fluorescein were recorded between 485 - 600 nm with an excitation at 460 nm. Similarly, the excitation spectra of the compounds were recorded in the 400 - 520 nm range with an emission at 520 nm. The experiments were conducted with an averaging time of 1 seconds; step of 0.5 nm; scan rate of 30 nm/min; Excitation slit width= 5 nm; Emission slit width = 5 nm; PMT Voltage=800 Volts

Fluorescence anisotropy *in-vitro* measurements were performed using the Cary 300 Fluorometer, with a set of two motorised polarisers, each of which is in front of the excitation source (a Xenon Lamp) and the detector, respectively. Four polarisation measurements were thus acquired based on the combination of the polariser orientations such as I_VV_, I_VH_, I_HH_, and I_HV_. Here H and V refer to horizontal and vertical orientations of excitation and emission polarisers, respectively.

Polarised intensity values for each compound were subtracted of the blank (buffer) polarisation intensity values and then the anisotropy is calculated using the equation:

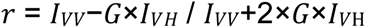

Calibration of the system was performed by measuring the fluorescence anisotropy of 10 nM fluorescein in glycerol solutions of different viscosities (w/w). The resulting anisotropy values were plotted as 1/r versus T/η (Perrin plot), where η is the viscosity of the solution and T was taken as 293 K. The slope of the Perrin’s plot reflects both the intrinsic fluorescence lifetime and the rotational diffusion time of a fluorescent species (*_τ_*_lifetime_/*_τ_*_rot_).

To determine compound binding to PDE4B, 5 nM fluorescein, MAN194, or AG3461 was incubated with increasing concentrations of purified PDE4B catalytic domain (0, 10, 20, 30, 100, 200, 300, and 500 nM; kind gift of Elisabetta Armani, Chiesi Farmaceutici, Italy) in the buffer described above. Samples were incubated for 60 min at room temperature before fluorescence anisotropy measurements were recorded. Binding curves were fitted using a single-site binding model in GraphPad Prism.

Specific binding was evaluated under three distinct conditions: in the first and second condition, free anisotropy of the compounds and the binding anisotropy of the compound in presence of 500 nM purified protein were recorded respectively. In the third condition, the purified protein was initially incubated with the unlabelled Rolipram (Sigma Aldrich) before adding the fluorescent-drug analogue to observe the resulting anisotropy. All the anisotropy experiments were conducted with an averaging time of 20 seconds; Excitation slit width= 5 nm; Emission slit width = 10 nm; PMT Voltage=800 Volts.

Fluorescence lifetime measurements of fluorescein, 8-F-cAMP, MAN194, and AG3457 were performed at a final concentration of 1 µM in TRIS buffer (pH 8). Measurements were carried out in a 10 mm quartz cuvette (Thorlabs) using a time-correlated single photon counting (TCSPC) setup (HORIBA FluoroMax, HORIBA Scientific). Excitation was provided by a pulsed diode laser (DeltaDiode, HORIBA Scientific) at 405 nm with a 100 MHz repetition rate and picosecond pulse width.

To obtain the instrument response function (IRF), 0.01% diluted Ludox in TRIS buffer (pH 8) was used. The detector photon counting sensitivity (α-value) was kept below 2% to avoid pile-up. Fluorescence decay data of the drug analogues were recorded under the same experimental conditions as the IRF. The decay curves were deconvoluted with the IRF using DA6 software (HORIBA Scientific). Depending on the sample, the data were fitted with single- or bi-exponential decay functions to obtain the fluorescence lifetimes.

### Plasmids, cell seeding and transfection

Plasmid DNA construct for PDE4B was subcloned into pVITRO2 with m-Ruby2 tag backbone using Gibson assembly kit (New England Biolabs). Site directed mutagenesis was performed to make the mutated mRuby plasmid. mCherry–Talin1 was cloned into the pEGFP-C1 backbone in place of EGFP.

### Glosensor assay

HEK293 cells were transfected using the GloSensor F22 plasmid (Promega) according to the protocol as described by (Brands et al. 2024). In Vial A, 50 µL of optiMEM was mixed with 1 µg of DNA, which included 25% GloSensor, 25% pcDNA and 50% PDE4B_mRuby plasmids. In a separate vial B, 50 µL of OptiMEM and 3 µL of PEI (Sigma Aldrich) (1 mg/mL) was mixed. The contents of both vials were mixed and incubated at room temperature for 10 minutes. In the meantime, cell suspension was created at 300k cells/mL. 1 mL of the cell suspension was added to the vial containing the mixture and then 100 µL of the cell-PEI-DNA suspension was loaded to each well of a Greiner 96 wells flat bottom plate. The above protocol would be sufficient for 8 wells in a 96 well plate and can be expanded as required. The cells containing plates were incubated for 48 hours in a cell culture incubator under optimal condition before performing the experiments.

On the day of the experiment, the cells were first washed with HBSS buffer (Capricorn Scientific) and 54 µL, 1 mM D-luciferin (MedChemExpress) in HBSS was added to the well. The cells were then incubated for 20 minutes at room temperature before recording the luminescence signals. Luminescence was recorded using CLARIOStar Plus (BMG Labtech) plate reader at 28°C. The measurements were done in kinetics mode for 24 cycles with a cycle time of 120 seconds. Emission was recorded between 580-80nm with a preset gain of 4000. Initial background was recorded for 3 measurement cycles before addition of the drugs. The drugs were previously serial diluted at 100x concentrations in 100% DMSO (Sigma Aldrich). Each of the concentrations were then re-diluted to 10x concentration from its corresponding 100x in HBSS buffer on the day of experiment. During the experiment 6 µL of the individual 10x concentrated drugs were added to the cells containing d-luciferin/HBSS mixture, which brought the final concentration of drugs to 1x in the cells. This helped to maintain constant DMSO concentration of 1% to the cells for all the different drug concentrations.

Data were analysed using Microsoft Excel and then plotted using GraphPad Prism. Each of the data points were initially corrected for background by dividing them with the average luminescence intensity of the measurements before drug addition. Then the data was normalised to the baseline (vehicle control – HBSS) and plotted as fold change in luminescence intensity over time for different concentrations. From the time series, the data points across different concentrations at 1920 seconds were used to plot the concentration response curve. The graph was fitted using the non-linear fit function in GraphPad Prism to obtain the IC50 values.

For the blocking assay, the cells were initially treated with 100 µM of MAN193 or vehicle control of MAN193 along with 1 mM D-luciferin for 30 minutes. The measurement conditions and setting were followed as described above. After recording the initial background of 3 cycles, IBMX (Sigma Aldrich) was added to the cells at final concentration of 30 µM or vehicle control. Data were corrected for background by dividing them with the average luminescence intensity of the measurements before drug addittion. Then the response induced by the vehicle of IBMX was subtracted from that obtained with IBMX and normalised the resulting values to the one of VC of MAN193 and plotted in the graph as time series. The area under the curve for the measurements was calculated and plotted using GraphPad Prism.

### Confocal microscopy

H9c2 cells were seeded to 0.01% PLL coated 8 well µ-Slide (Ibidi GmbH) at 15000 cells per well. Experiments were performed 24 hours after the cells were seeded so as to allow the cells to attach and spread properly. On the day of the experiment, the cells were treated with 10 nM MitoTracker Deep Red (Thermo Fischer Scientific) for 15 minutes and then washed once with LCIM. The cells were then incubated with 100 µM rolipram or 2% DMSO for 30 minutes. Molecular brightness measurements were performed using a Leica SP8 confocal microscope. The confocal images were obtained as 1024 × 1024 pixels with a pixel size of 100 nm. A time series was recorded for 60-70 frames with a frame interval of 20 seconds (Işbilir et al. 2021). The fluorescein emission was imaged using the 488 nm laser line with 2% power and emission was detected between 500-550 nm. Furthermore, MitoTracker deep red was imaged using the 633-laser line with 2% power and emission was detected between 650-750 nm. Once the acquisition started, fluorescent drug analogue was added to the cells (final concentration in cell 10 µM) after 5 initial frames. The confocal micrographs were pre-processed as shown in **Supplementary Figure 6** before measuring the brightness values. The MitoTracker images were converted into mask using the “Convert to Mask” feature in ImageJ. The masked images were then subtracted from the fluorescein emission channel images and saved in tiff format. ROIs arising from cells were generated using the Cellpose TrackMate plugin and all the ROIs generated from an individual cells were saved as a single file. The micrographs and ROI files were then processed using a custom written ImageJ Fiji macro. The macro calculated the variance and mean intensity values and gave the output as molecular brightness across the time series, which is the ratio of variance to mean.

Molecular brightness measurements were also performed in cell over expressing PDE4B. 24 hours after the cells were seeded, they were transfected using Lipofectamine 2000. 1 µg of plasmid DNA (PDE4B_mRuby), 2 µL of Lipofectamine 2000 and 150 µL of OptiMEM were mixed and incubated at room temperature for 15 minutes and then added in equal portions to the cells. Twenty-four hours after transfection, the culture medium was replaced with fresh medium, and cells were incubated for a further 24 h prior to imaging. On the day of the experiment the cells were treated as described previously. Fluorescence lifetime measurements in cells were performed using the Leica Stellaris 8 system. The data was analysed using FilmJ plugin.

### Confocal microscopy of focal adhesions

H9c2 cells were seeded on fibronectin coated 8 well µ-Slide (Ibidi GmbH) at 7000 cells per well. 24 hours after cell seeding, the cells were transfected using Lipofectamine 2000. 1 µg of plasmid DNA (50% mCherry Tln1 and 50% PDE4B), 2 µL of Lipofectamine 2000 and 150 µL of OptiMEM were mixed and incubated at room temperature for 15 minutes and then added in equal portions to the cells. After 24 hours the cell media were replaced with fresh media and more 24 hours was waited for the experiments. On the day of the experiment, the cells were either incubated with 100 µM rolipram or 2% DMSO containing live cell imaging media for 30 minutes and then treated with 10 µM of MAN193 for another 30 minutes before imaging. Localisation microscopy was performed using a Zeiss LSM800 confocal microscope. The confocal images were obtained as 1024 × 1024 pixels with a line averaging of 2. mCherry-Talin1 was imaged using the 561-laser line with 4% power and emission was detected between 600-650 nm. Furthermore, the MAN193 emission was imaged using the 488 nm laser line with 0.5% power and emission was detected between 500-550 nm. Once the images were obtained, they were processed using ImageJ.

### Fluorescence Anisotropy Imaging

Anisotropy imaging was performed using a custom-built setup as described in **Supplementary Figure 6**. The microscope is built on a Leica, DMIRE2 body and uses a 488 nm diode laser (OBIS, Coherent) for excitation. Once the excitation light is separated from the emission using a dichroic mirror (ZT 405/488/635, Chroma) the beam is circularly polarised using a quarter wave plate and spatially filtered using a confocal system and then reflected on the back focal plane of the microscope. Before the excitation beam enters the microscope, it is either horizontally or vertically polarised using a polarised beam splitter. Horizontal and vertical polarisation are defined with respect to the forward moving plane of the excitation beam. Once the emission beam is emitted by the sample, it is filtered using a 572/26nm emission filter and transmitted through the beamsplitter Optosplit II (Cairn Technology). Once the beams are split, they are further filtered using two linear polarisers placed in horizontal and vertical polarisation positions. The split beams are collected on an EM-CCD (Andor, iXON DV897). This allows simultaneous imaging of both polarisation vectors and enables all the four polarisation vectors to be obtained as required for generation of G-factor images.

Fluorescence images of 100 nM fluorescein in 50% glycerol and water were obtained in all the four polarisation vectors by rotating the polarised beam splitter as described in the **Supplementary Figure 8** to calculate G-factor. Once the mean intensities were obtained from each image using ImageJ Fiji, the mean intensities were subtracted from the mean intensities of LICM along the polarisation vectors and g-factor was calculated. The g-factor is calculated during each experimental session to account for small variations in the optical setup alignment.

For the anisotropy imaging experiments, HEK293 cells were seeded in #1.5 glass cover slips at a density of 250,000 cells and allowed to grow for 18-24 hours before the experiments. On the day of the experiments, the cells were either incubated with 100 µM rolipram or 2% DMSO containing live cell imaging media for 30 minutes and then treated with 100 nM of MAN193 or 50 nM Fluorescein diacetate for another 30 minutes. The cells were then washed once with live cell imaging media, and the cover slips were loaded into a Attofluor Cell Chamber (Thermo scientific) with 300 µL of LICM to prevent the cells from drying during the experiments. The cells were imaged using a 40x objective at 512 x 512 pixels and 50 frame movies were recorded with 5 second intervals. The images were acquired using exposure of 100 ms and an EM gain of 200. The EMCCD was always electronically cooled to −70°C while recording the measurements to avoid accumulation of environmental noises.

Once the two polarisation images containing movies were obtained, using a custom written ImageJ macro, the corresponding anisotropy movie was generated with the calculated g-factor. The anisotropy movies were then processed to obtain the modal anisotropy values from each frame of the movies. Using a custom imageJ macro the anisotropy histogram from each of the frames were obtained. Histogram here represented the distribution of anisotropy values across the 256 bins of the image and its corresponding frequencies. A running average was run across the whole data using ±2 data points to smooth the data. The maximum modal anisotropy values from this data were obtained using the following formula in Microsoft excel: =INDEX($A:$A, MATCH(MAX(C1:C256), C:C, 0))

The modal anisotropy values from the first 10 frames were averaged to obtain the quantitative anisotropy values for each of the experimental conditions. The values were plotted and tested for significance using Anova test using GraphPad Prism.

